# Multiplexed CRISPR/Cas9 mutagenesis of rice PSBS1 non-coding sequences for transgene-free overexpression

**DOI:** 10.1101/2023.10.20.563333

**Authors:** Dhruv Patel-Tupper, Armen Kelikian, Anna Leipertz, Nina Maryn, Michelle Tjahjadi, Nicholas G. Karavolias, Myeong-Je Cho, Krishna K. Niyogi

## Abstract

Understanding CRISPR/Cas9’s capacity to generate native overexpression (OX) alleles would accelerate agronomic gains achievable by gene editing. To generate OX alleles with increased RNA and protein abundance, we leveraged multiplexed CRISPR/Cas9 mutagenesis of non-coding DNA sequences located upstream of the rice *PSBS1* gene. We isolated 120 transgene-free, gene-edited alleles with varying NPQ capacity *in vivo* —ranging from complete knockout to overexpression, using a high-throughput phenotyping and transgene screening pipeline. Overexpression of *OsPSBS1* increased protein abundance 2-3-fold, matching fold changes obtained by transgenesis. Increased PsbS protein abundance enhanced non-photochemical quenching capacity and improved water-use efficiency. Across our resolved genetic variation, we identify the role of 5’UTR indels and inversions in driving knockout/knockdown and overexpression phenotypes, respectively. Complex structural variants, such as the 252kb duplication/inversion generated in this study, evidence the potential of CRISPR/Cas9 to facilitate significant genomic changes with negligible off-target transcriptomic perturbations. Our results may inform future gene-editing strategies for hypermorphic alleles and have opened the door to the pursuit of gene-edited, non-transgenic rice plants with accelerated relaxation of photoprotection.

## Introduction

Optimizing photosynthetic efficiency is one of the most promising routes to engineering more sustainable and productive crop varieties^1,2^. Several recent breakthroughs have showcased the use of informed photosynthetic design principles to improve agronomic traits in the field. A photorespiratory bypass, made possible in part by cross-phyla overexpression of green algal glycolate dehydrogenase, increased field-grown biomass of *Nicotiana tabacum* by 20%^3^ and enhanced relative fitness over the wild type (WT) when grown under elevated temperatures^4^. Overexpression of putative transcriptional regulators OsDREB1C^5^ and zmm28^6^ have also shown significant agronomic yield gains in model rice and elite maize varieties, respectively, although the causal effects on photosynthesis are still unclear. Transgenic overexpression of three highly conserved genes from *Arabidopsis thaliana* (*AtVDE*, *AtPsbS*, and *AtZEP*, hereafter VPZ), which are involved in photoprotection through non-photochemical quenching (NPQ), increased relaxation of but did not compromise NPQ under fluctuating light conditions, resulting in increases in tobacco biomass by 15%^7^ and soybean seed yield by up to 20%^8^ in small-scale field trials. The prospects for rationally designing photosynthesis in agronomic environments are bright.

However, these approaches relied on expression of foreign DNA, or transgenes, which incur regulatory complications and can be susceptible to gene silencing across generations^9^. NPQ genes are found in all plants, and NPQ proteins are highly conserved in their function. We reasoned that altering endogenous gene expression could achieve similar agronomic gains without the need for persistent transgenes. As a proof of concept, we targeted the rice Photosystem II Subunit S (*OsPSBS1*) gene, a core factor in high-light and fluctuating-light tolerance, to generate mutants with increased *OsPSBS1* expression and improved NPQ capacity.

Quantitative variation in rice *OsPSBS1* has already been observed between *japonica* (higher NPQ) and *indica* (lower NPQ) subspecies^10^, potentially highlighting NPQ as a trait that has recently undergone selection. Transgenic overexpression of PsbS in rice has been shown to increase radiation-use efficiency in greenhouse-grown plants^11^, and overexpression in tobacco has been implicated in increasing intrinsic water use efficiency (iWUE) under red light^12^. Rice is a diploid model C_3_ monocot with single-copy orthologs encoding VDE, ZEP, and a single, functional PsbS (OsPsbS1)^13^. And as a food crop that accounts for over 20% of all global calories consumed, rice is an impactful and genetically tractable target for improvement.

CRISPR/Cas9 has dramatically expanded our capacity to produce targeted loss-of-function mutations. Recently, CRISPR/Cas9 editing of cis-regulatory elements (CREs) has been used to decrease, rather than abolish, gene expression in tomato^14^ and maize^15^, improving fruit-and grain-yield related traits. Similar multiplexed strategies have shown the potential of knocking out known CREs in pathogen susceptibility^16,17^, or in identifying and editing novel CREs that constrain tradeoffs between rice grain yield and plant architecture^18^. Though newer toolkits such as the CRISPR-Cas12a promoter editing (CAPE) system^19^ and approaches such as promoter swapping^20^ show promise in advancing quantitative trait engineering efforts, finer understanding of ways to significantly increase gene expression without the use of persistent transgenes is still lacking.

To address these challenges, we established a high-throughput pipeline for screening novel alleles of *OsPSBS1* generated by CRISPR/Cas9 mutagenesis of upstream non-coding sequences (NCS). We generated a mutant library covering a large phenotypic range of *OsPSBS1* gene expression, including overexpression, with significant effects on NPQ and iWUE. Finally, we identify themes in cis-regulation across our library that may inform gene editing strategies for knockdown or overexpression of genes that affect other desirable plant traits.

## Results

### High-throughput screening of edited, semi-dominant OsPSBS1 promoter alleles

To generate novel cis-regulatory variation upstream of *OsPSBS1*, a CRISPR/Cas9 construct targeting eight specific and conserved rice gRNA sites (**Supplementary Table 1**) was introduced into Nipponbare (sp. *japonica*) rice calli via *Agrobacterium*-mediated transformation. The designed gRNA target regions distal (green triangles) and proximal (magenta triangles) to the *OsPSBS1 gene,* avoiding a putative QTL for NPQ activity^10,13^ (**Fig. 1a**). 23 fertile, independent transformants were generated, and multiple plant lines were recovered from each transformed callus, when possible.

**Figure 1.**
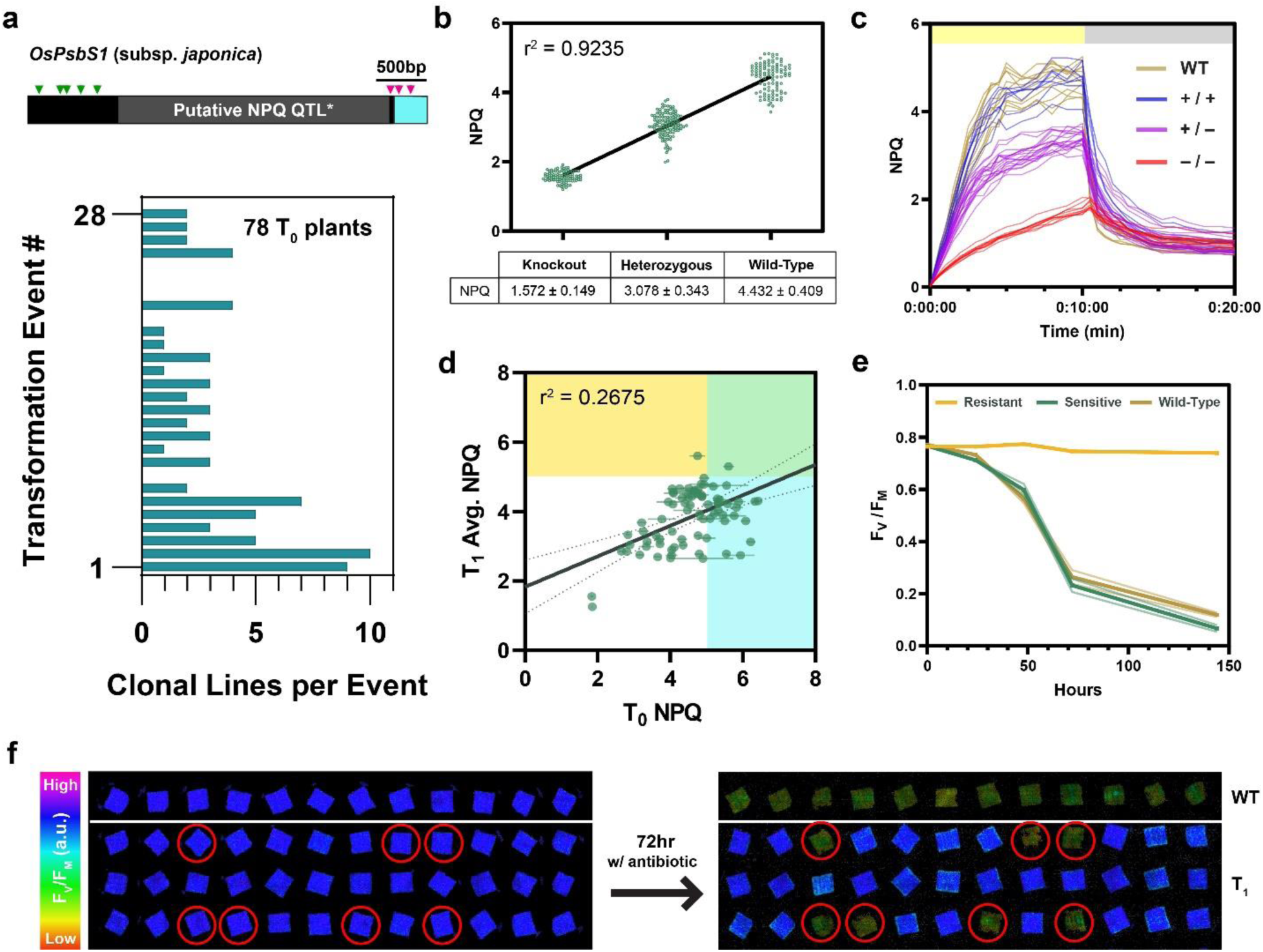
A high-throughput chlorophyll fluorescence to resolve T_1_ quantitative variation and Cas9 transgene segregation. **(a)** Target gRNA sites (triangles) distributed in distal (green) and proximal (magenta) regions upstream of *OsPSBS1*, with an insertion specific to subsp. *japonica* varieties marked in gray^1,2^. Multiple clonal lines were regenerated across 23 independent events. **(b)** Linear regression of maximum NPQ capacity after 10 min of 1500 µmol photons m^-2^ s^-1^ of blue light in PsbS knockout, heterozygous, and WT lines (n = 95, 117, and 101 biological replicates respectively). **(c)** Chlorophyll fluorescence phenotyping of NPQ resolves homozygous alleles (+/+, blue; -/-, red) in 36 progeny of a single segregating T_0_ parent relative to WT (n=12, brown), all replicates shown. **(d)** Correlation of T_0_ max NPQ with the average max NPQ of its corresponding T_1_ alleles (n = 78 T_0_ plants, 2-4 biological replicates per T_1_ population). Individuals with NPQ exceeding +2 SD of WT in the T_0_ generation, T_1_ generation, or both generations are shown in cyan, yellow, and green respectively. **(e,f)** Addition of the plant selection antibiotic hygromycin to leaf punches phenotyped in (c) identifies sensitive individuals with inhibited F_v_/F_m_ (that lack the Cas9 transgene (circled).

High-throughput *in vivo* chlorophyll fluorescence screening was used to resolve stable, heritable phenotypes across the 78 diploid T_0_ transformants, yielding up to 156 gene-edited alleles. Our approach leveraged the fact that PsbS is semi-dominant and shows a strong linear correlation between gene copy number and NPQ capacity (r^2^ = 0.9235) (**Fig. 1b**). Thus, in a segregating population, it is possible to resolve individuals with divergent, homozygous alleles by phenotype alone (**Fig. 1c**). This was important as phenotypes in the T_0_ generation may be somatic, and competition between alleles may mask changes in gene expression and activity. Notably, maximum NPQ in the T_0_ generation was a poor predictor of stable T_1_ phenotypes (r^2^ = 0.2675). T_0_ phenotypes overestimated the number of putative T_0_ overexpressors (blue) and failed to identify a candidate stable T_1_ overexpressor line (yellow) (**Fig. 1d**).

This screen for NPQ phenotypes was extended to incorporate segregation of the hemizygous Cas9 transgene. Leaf punches used to assess NPQ capacity (**Fig. 1c**) were treated with hygromycin, the antibiotic used for selection of transformants, and monitored for a decline in the quantum efficiency of photosystem II (F_v_/F_m_), a common indicator of plant stress (**Fig. 1e,f**). Obvious differences in resistance could be observed within 72 h of incubation, with a decrease in F_v_/F_m_ of over 75% upon loss of Cas9, as verified by PCR (**Supplementary Fig. 1**). By screening pools of 36-72 T_1_ progeny per T_0_ parent, it was possible to isolate multiple individuals harboring homozygous gene-edited alleles and lacking Cas9 by Mendelian segregation of each trait, allowing us to rapidly identify fixed germplasm for further analysis in the T_2_ generation.

### Gene-edited, overexpression alleles are present, albeit rare

Putative homozygous alleles were identified via pairwise comparison between WT plants and progeny from a single T_0_ parent, as depicted in **Fig. 1c**. To assess the variation in phenotypes across all alleles, maximum NPQ for all 120 homozygous alleles was plotted in **Fig. 2**.

**Figure 2.**
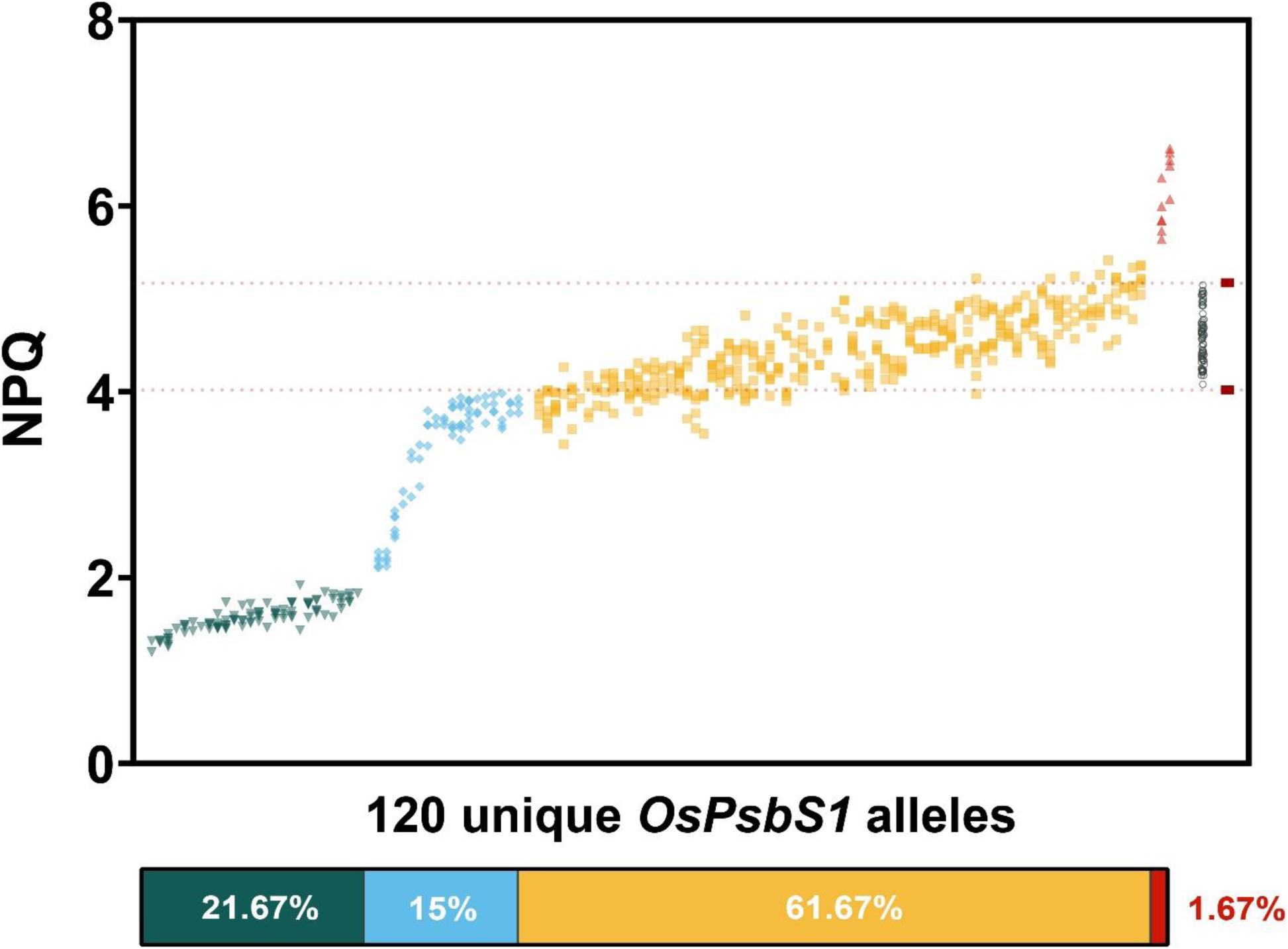
Maximum NPQ of 120 homozygous, gene-edited alleles spanning 78 T_0_ events. Max NPQ (n=1-4 technical replicates each, 2-8 biological replicates each) after 10 minutes at 1500 µmol photons m^-2^ s^-1^ blue light exposure. Each independent allele is sorted from lowest to highest average NPQ capacity. WT NPQ of 64 biological replicates spanning all phenotyping experiments is shown (gray, open circles on the right) with boundaries demarcating ±2 SD (red, dashed lines). Lines with all biological replicates below/above the WT boundary were binned by phenotype: knockout (teal, inverse triangle), knockdown (blue, diamond), WT-like (yellow, square), and overexpressor (red, triangle). Proportions of each observed phenotype are shown below.

Almost two-thirds of the 120 stable alleles isolated were WT-like in NPQ capacity (61.67%), with the second and third largest groups being knockout (KO, 21.67%) and knockdown (KD, 15%) phenotypes, respectively. Two independent overexpression alleles (OX) were isolated, comprising 1.67% of the total phenotypic variation.

### Representative alleles confirm phenotype-by-expression relationships

A panel of representative alleles spanning the knockout to overexpression spectrum was selected to correlate observed NPQ phenotypes with gene expression and protein abundance. Interestingly, the full range of phenotypic diversity was present among clonal lines derived from a single transformation event (Event 2), with the only other overexpression allele arising from Event 19. Notably, Event 2 also produced the greatest number of regenerated lines (**Fig. 1a**). This observation demonstrates that most gene editing occurred after initial selection of transformation events, and sister lines can experience divergent Cas9 editing trajectories.

Greenhouse-grown plants were sampled 1 h after sunrise and 1 h before sunset and analyzed for differences in *OsPSBS1* transcript levels. The Event 2-5 KO line (hereafter referenced using the nomenclature “Event # - Line # _ Phenotype”) was a useful internal control, because unlike most other phenotypic KO alleles generated, 2-5_KO contains a large deletion that spanned the entire *OsPSBS1* coding sequence. Correspondingly, this allele had gene expression well below the threshold of detection by our *OsPSBS1*-specific primers (Log_2_FC < -10). We observed some stochasticity in gene expression across experiments; for example, there was a significant decrease in gene expression in 2-6_KD in the morning (**Fig. 3a**) but not the evening (**Fig. 3b**). Conversely, we found a statistically significant 4-8 fold increase in *OsPSBS1* transcript levels in the evening dataset (**Fig. 3b**) but not the morning dataset, although trends in KD or OX lines across datasets were largely consistent.

**Figure 3.**
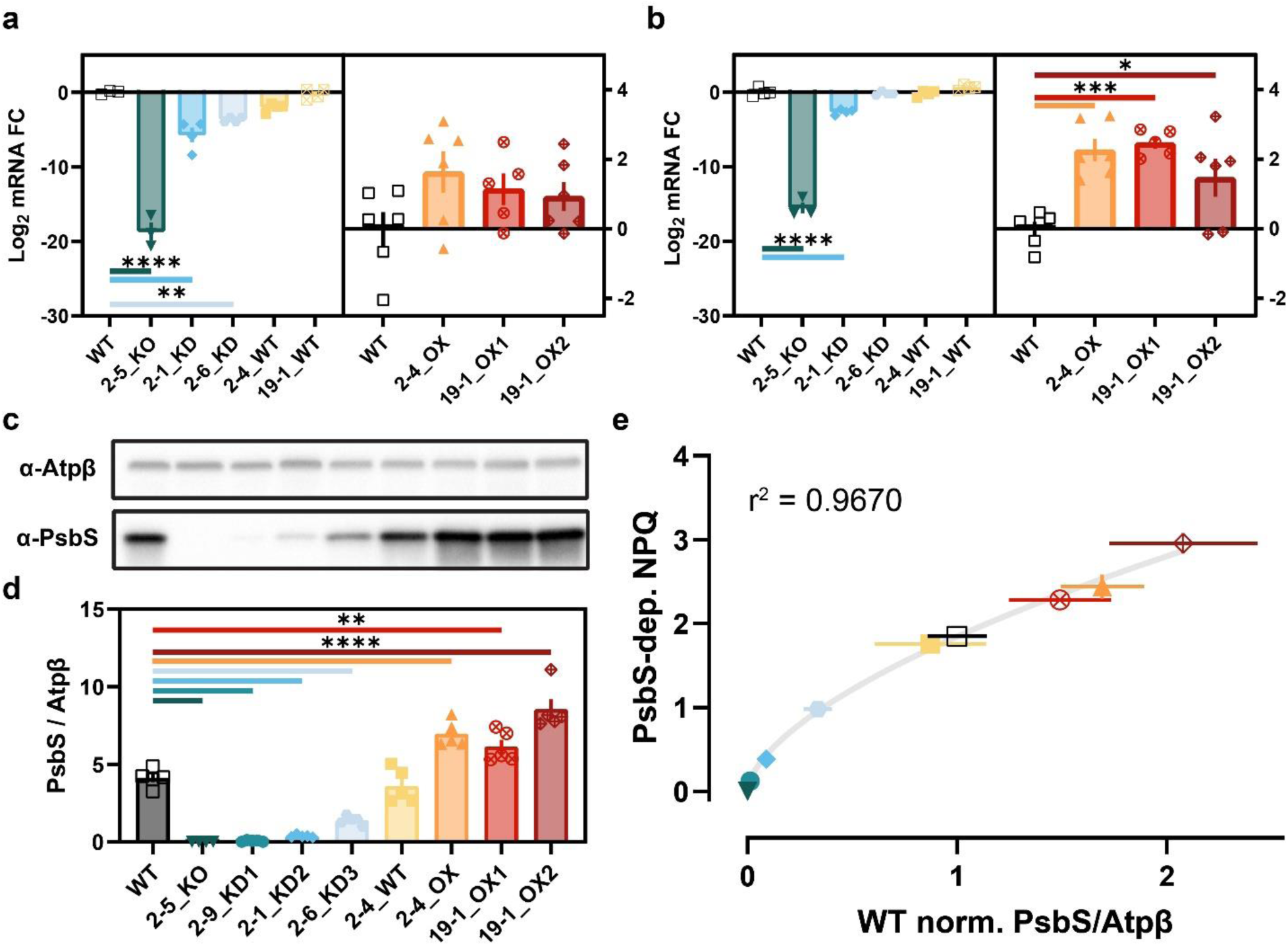
OsPSBS1 transcript expression and protein abundance across 8 representative alleles. Log_2_ relative *OsPSBS1* transcript quantification against WT, normalized to *OsUBQ* and *OsUBQ5*. Samples were collected across **(a)** morning and **(b)** evening experiments (n=3-6 biological replicates, data ± SEM). Gene-edited alleles are ordered from lowest to highest total NPQ. **(c)** Representative immunoblots (6 µg total protein) of OsPsbS1 and the chloroplast loading control Atpβ. **(d)** Normalized OsPsbS1 immunoblot band intensity shown ± SEM across 4 (2-5_KO) or 5 (remaining genotypes) individual replicates each. **(e)** PsbS-dependent NPQ capacity calculated by determining steady-state NPQ at 2000 µmol photons m^-2^ s^-1^ subtracted from the average residual NPQ in the 2-5_KO line. NPQ is plotted against WT-normalized OsPsbS1/Atpβ band intensity (mean ± SEM shown). A logarithmic curve fit is shown in gray. For all panels, genotypes and replicates are shown: WT (black, open square), 2-5_KO (dark teal, inverse triangle), 2-9_KD1 (teal, circle), 2-1_KD2 (blue, diamond), 2-6_KD3 (light blue, hexagon), 2- 4_WT (yellow, square), 2-4_OX (orange, triangle), 19-1_OX1 (red, crossed circle), 19-1_OX2 (maroon, crossed diamond). Pairwise significance was determined by ordinary one-way ANOVA (α =0.05) using Dunnett’s test for multiple comparisons against Nipponbare WT (*p≤0.05, **p≤0.01, ***p≤0.001, ****p<0.0001).

OsPsbS1 protein abundance was determined by immunoblot, and a set of representative experiments are shown in **Fig. 3c**. Quantification of normalized band intensity across replicates confirmed protein abundances spanning 1.5% to 250% of WT OsPsbS1 levels (**Fig. 3d**), with all blots reported in **Supplementary Fig. 2**. The observed logarithmic best-fit between protein abundance and PsbS-dependent NPQ (**Fig. 3e**) mirrors what has been reported in Arabidopsis^21^.

### Knockout and knockdown alleles arise from variation in the 5’UTR

To assess the causal mutations underlying these phenotypes, the approximately 4.3-kb upstream cis-regulatory region was amplified and sequenced by Sanger sequencing. 107 of the 120 unique alleles (**Fig. 2**) could be amplified by PCR (89.2%). Several alleles containing large deletions of the five distal gRNA target sites (**Fig. 1a**) were detected. Interestingly, these lines had NPQ capacity that was indistinguishable from WT (**Supplementary Fig. 3**). Thus, most of the observed phenotypic variation can be explained by variants within and near the 5’UTR.

We utilized mVista^22^ to assess variation in conserved non-coding sequences (CNS) proximal to *OsPSBS1* across 10 representative species. We found no significant conservation across five dicot genome comparisons but saw consistent, high-confidence (> 50%) conservation within the 5’UTRs of the *PSBS* gene in five representative monocot genomes (**Fig. 4a**).

**Figure 4.**
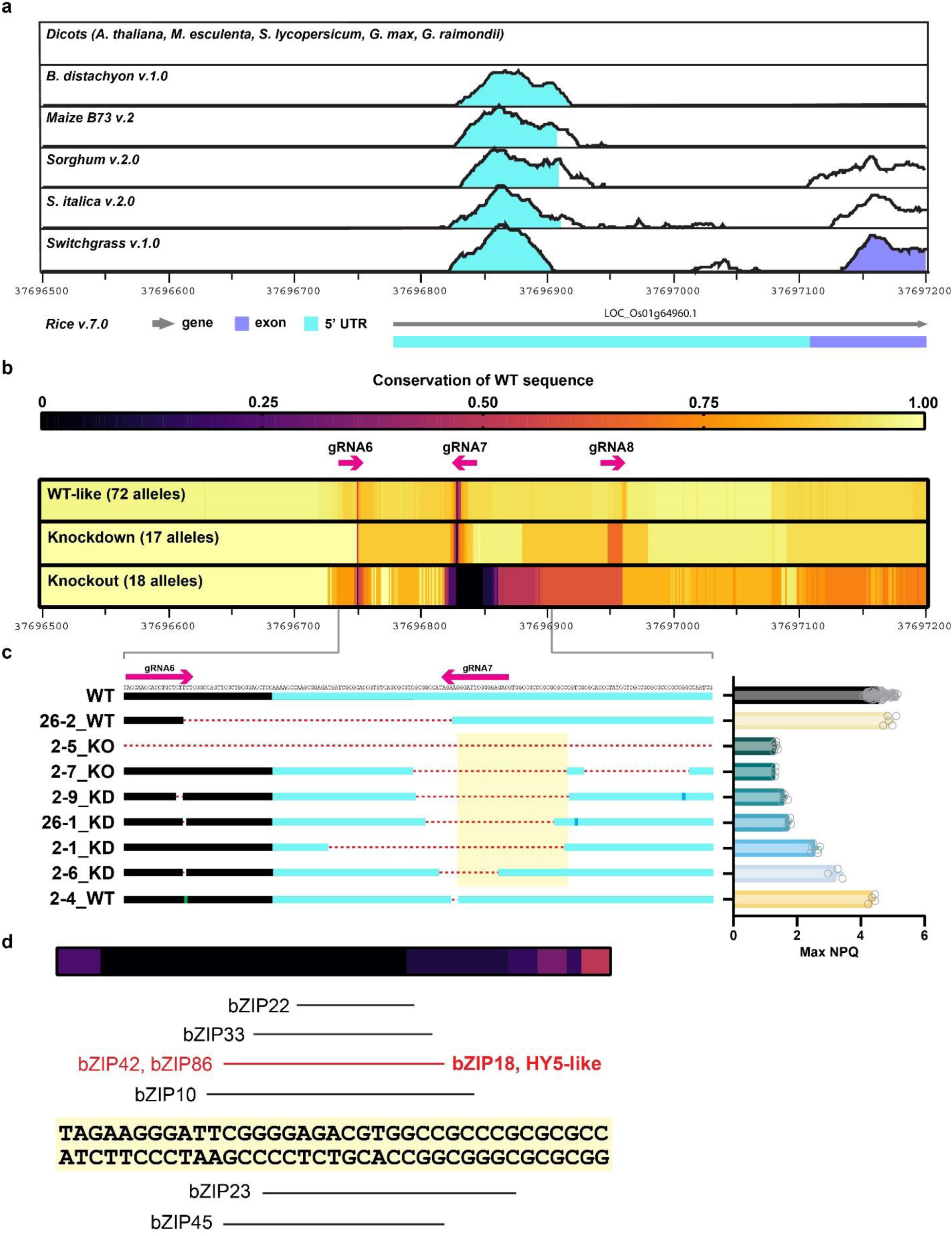
Aggregate analysis of gene-edited knockout and knockdown alleles. **(a)** mVista alignments of the proximal OsPsbS1 genomic locus [Chr01: 37,696,500 – 37,697,200] against 5 dicot and 5 monocot species. Shading indicates conservation exceeding 70% within the 5’ untranslated region (UTR, cyan) or coding sequence (exon, purple) of the *OsPSBS1* gene. **(b)** Conservation of WT sequence across gene-edited variants within the proximal *OsPSBS1* region, aggregated by phenotype. Target sites and directionality of gRNA are shown as magenta arrows. **(c)** A zoomed-in subset of representative KO and KD lines with small deletions spanning sites with low sequence conservation. Max NPQ phenotypes from (Fig. 2) are shown to the right of each allele (n=3-5 biological replicates each, excluding WT). The yellow highlighted region in **(c)** is shown at higher nucleotide resolution in **(d)** relative to aggregate WT sequence conservation across KO alleles. Putative transcription factor binding sites predicted by PlantRegMap^3^ are shown.

Using our library of sequenced *OsPSBS1* non-coding sequence alleles, we assembled a phenotype-aggregated heatmap to correlate indels at the three proximal gRNA (gRNA 6-8) sites with phenotype (**Fig. 4b**). As expected, mutations are found most frequently at the 3’ NGG-proximal end of each gRNA. WT-like alleles showed that small indels can be tolerated without significant phenotypic cost. In contrast, shared large deletions within the monocot-conserved CNS were associated with KO phenotypes. KD alleles largely maintained the WT sequence within the CNS, but they had larger, variable indels (50-70% conservation) nearby that reduced but did not abolish *OsPSBS1* gene expression.

Several representative alleles with relatively small deletions were disaggregated to better understand the causal locus for *OsPSBS1* expression (**Fig. 4c**). Interestingly, the transcription start site (TSS) appears to be completely dispensable and is not required for WT NPQ capacity. In contrast, deletions proximal to gRNA7 are associated with a range of KO and KD phenotypes, dependent on deletion size and location. We further interrogated the highlighted region in Fig. 4c, which encompassed the region of poor WT sequence conservation across KO lines (0-11%). Notably, this locus contains several high confidence (p < 0.0001) binding sites for bZIP transcription factors^23^, including bZIP18, a HY5-like ortholog (**Fig. 4d**).

### Complex structural variants underlie overexpression

Unexpectedly, neither OX allele could be resolved by PCR genotyping. To determine whether complex, structural variants were underlying these phenotypes, a homozygous 2-4_OX T_2_ line and segregating 19-1_OX T_1_ line were sequenced by long-read HiFi circular consensus sequencing (Pacific Biosciences). Reads were assembled *de novo*, mapped onto the Nipponbare *O. sativa* v7.0 reference genome^24^, and used to produce dot plots showing sequence and structural variants (**Supplementary Fig. 4a,b**).

Zooming in on the *OsPSBS1* locus on Chromosome 1 revealed structural variation that was not observable at the whole-genome scale. The 2-4_OX line showed robust signatures of a ∼252-kb inversion via dot plot (**Supplementary Fig. 5a**), which was further substantiated at the sequence level by looped, head-to-head single long-read sequences visualized by Integrative Genomics Viewer (IGV)^25^ (**Supplementary Fig. 5b**). The increased read depth, confirmed by quantitative PCR of genomic DNA copy number (**Supplementary Fig. 5c**), verified that the 2- 4_OX allele was a 252-kb duplication/inversion.

In contrast, the segregating 19-1_OX line showed no appreciable differences by dot plot (**Supplementary Fig. 6a**). However, local inspection revealed the presence of a smaller ∼4-kb inversion with a Sniffles variant-called allele frequency of 0.361 (**Supplementary Fig. 6b**), which was successfully fixed to homozygosity (19-1_OX1). Despite this small genomic perturbation, we saw significant decreases in aboveground and grain biomass in a subset of dwarfed 19-1_OX progeny, hereafter referred to as 19-1_OX2 (**Supplementary Fig. 7a**). This was not true of the fixed 19-1_OX1 or 2-4_OX allele, which grew similarly to azygous WT controls (**Supplementary Fig. 7b**). Notably, isolating transgene-free 19-1 lines was confounded by the presence of 3 T-DNA insertions, as inferred from Mendelian segregation of antibiotic sensitivity (**Supplementary Fig. 7c**). Additionally, while the 19-1_WT (Cas9^-^) and 19-1_OX1 (Cas9^+^) had 100% heritable alleles, 19-1_OX2 could not be fixed to phenotypic homozygosity even in the T_4_ generation (**Supplementary Fig. 7d**). It is unclear if a linked, repetitive element found at low frequency by Pacbio sequencing may be involved in the dwarf or NPQ OX phenotypes (**Supplementary Fig. 6b**), but the presence of dwarfed plants with WT NPQ capacity suggests this phenotype is independent of the gain-of-function mutation. In summary, both stable *OsPSBS1* overexpression alleles (2-4_OX and 19-1_OX1) involve inversions upstream of the gene (**Fig. 5**).

**Figure 5.**
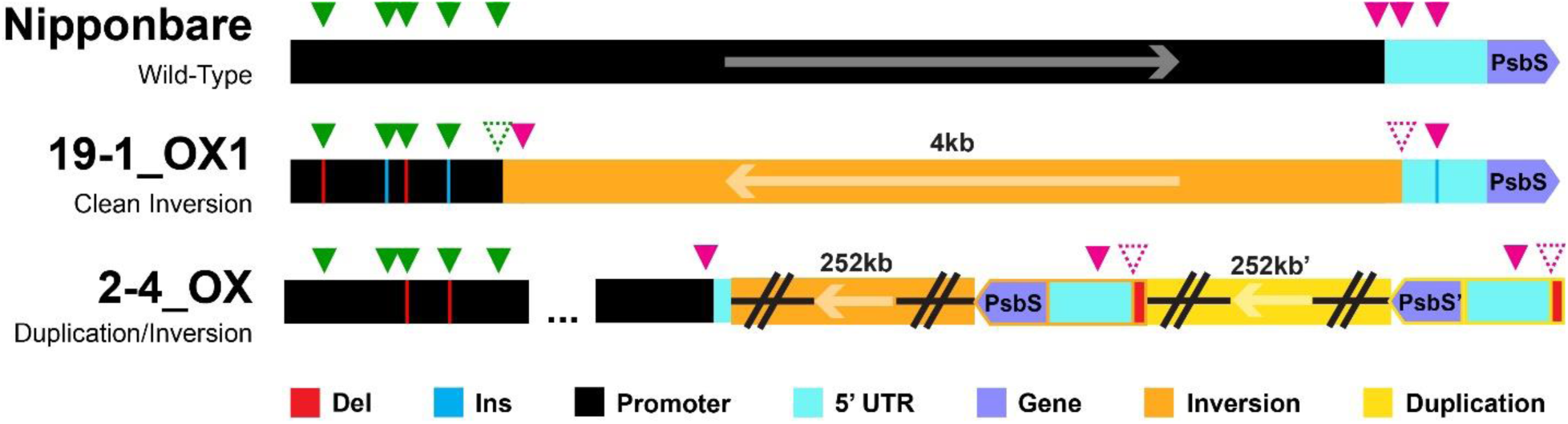
Long-read, whole-genome sequenced alleles driving OsPsbS1 overexpression. Cartoon depictions of the stable 2-4_OX and 19-1_OX1 overexpression alleles relative to the WT promoter. Distal gRNA sites are marked by green triangles, proximal gRNA sites by magenta triangles, the 5’UTR in cyan, and the *OsPSBS1* gene in purple. 1-3bp deletions (red lines) and insertions (blue lines) are shown relative to inversions (orange) and/or duplications (yellow). Structural variants that disrupt gRNA binding sites are shown with dashed triangles.

### Transcriptomic analysis of the duplication/inversion

The 252-kb duplication/inversion was a sizeable, drastic genomic change generated by CRISPR/Cas9 mutagenesis. To determine the broader consequences of such a structural change, we performed an RNAseq experiment to assess the extent of differential gene expression between the 2-4_OX and 2-4_WT lines. In total, 104 differentially expressed genes (DEGs) were resolved using a generous adjusted p-value (≤ 0.1), representing less than 0.2% of all genes within the genome. 22 of all the significantly DEGs were contained within the duplication/inversion, representing 21.2% of all DEGs and 62.9% of all genes within the duplication/inversion (**Fig. 6a**).

**Figure 6.**
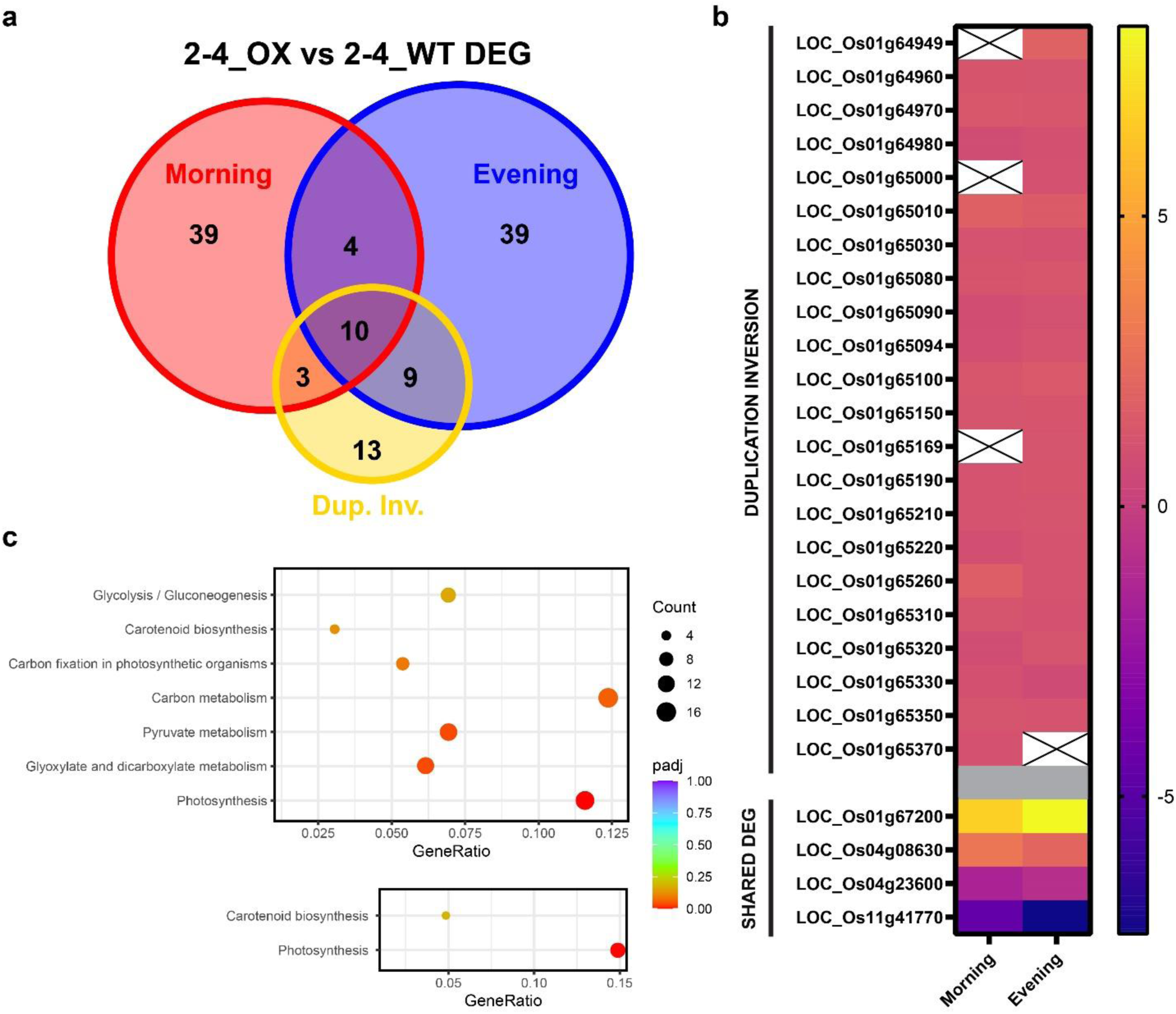
Transcriptome analysis of the 2-4_OX duplication/inversion. **(a)** Number of shared and unique differentially expressed genes (DEGs, adjusted p-value ≤ 0.1) between 2-4_OX and 2-4_WT alleles across morning (red) and evening (blue) samples relative to the 35 genes within the 252 kb duplication/inversion (yellow). **(b)** Log_2_ fold change gene expression of DEGs within (top) or outside (bottom) the duplication inversion across morning and evening datasets. **(c)** KEGG pathway enrichment (adj. pval ≤ 0.2) of all DEG (p-value < 0.05) within the morning dataset (top) or across both morning and evening datasets (bottom).

Next, we assessed the log_2_ fold changes of individual DEGs. We observed a consistent 2-fold increase in expression (Average Log2FC = 1.123 ± 0.255 SD) across the 22 differentially expressed duplication/inversion genes within either or both datasets. Of the remaining DEGs, only 4 were shared across both datasets, including the strongest upregulated DEG: a ribosomal L13 family protein that was negligibly expressed in the 2-4_WT allele but actively expressed in the 2-4_OX allele (**Fig. 6b**). We leveraged KEGG pathway enrichment to infer broader physiological consequences due to DEG upregulation of *OsPSBS1* and nearby genes by the duplication/inversion. Of these pathways, photosynthesis and carbon assimilation-related metabolic pathways were significantly enriched (**Fig. 6c**).

### Varying PsbS abundance impacts red light-dependent gas exchange phenotypes

Previous work has shown that transgenic overexpression of PsbS does not compromise steady-state CO_2_ assimilation in rice^11,26^, but in tobacco it can increase maximum intrinsic water-use efficiency (iWUE, A_n_/g_sw_) under red light^12^. To determine the extent to which these phenotypes are consistent in rice, seven phenotypically diverse genotypes were assessed for steady-state chlorophyll fluorescence and gas exchange phenotypes under increasing red light relative. This required removing the contribution of stress-related effects on iWUE independent of plant genotype, constraining the minimum F_v_/F_m_ threshold of analyzed replicates to 0.803 (**Supplementary Figure 8a**).

Knockdown lines 2-9_KD1, 2-1_KD2, and 2-6_KD3 showed significantly lower NPQ (p < 0.0001), and overexpression lines 19-1_OX1 and 19-1_OX2 had significantly higher NPQ (p < 0.0001), at all light intensities greater than 500 µmol photons m^-2^ s^-1^ relative to the WT control. In contrast, 2-4_OX showed significantly higher NPQ only at 1500 and 2000 µmol photons m^-2^ s^-1^ (p<0.0001) (**Fig. 7a**). Concurrent with the expanded higher NPQ capacity range, the 19-1_OX1 and 19-1_OX2 alleles exhibited a significantly lower ΦPSII (**Fig. 7b**), CO_2_ assimilation rate (A_n_, **Fig. 7c**), and stomatal conductance (g_sw_, **Fig. 7d**) compared to WT. The remaining Event 2 genotypes showed little significant difference across ΦPSII, A_n_, or g_sw_, excluding a modest increase (p < 0.05) in CO_2_ assimilation rate in 2-5_KO at light intensities over 1200 µmol photons m^-2^ s^-1^ (**Fig. 7c**).

**Figure 7.**
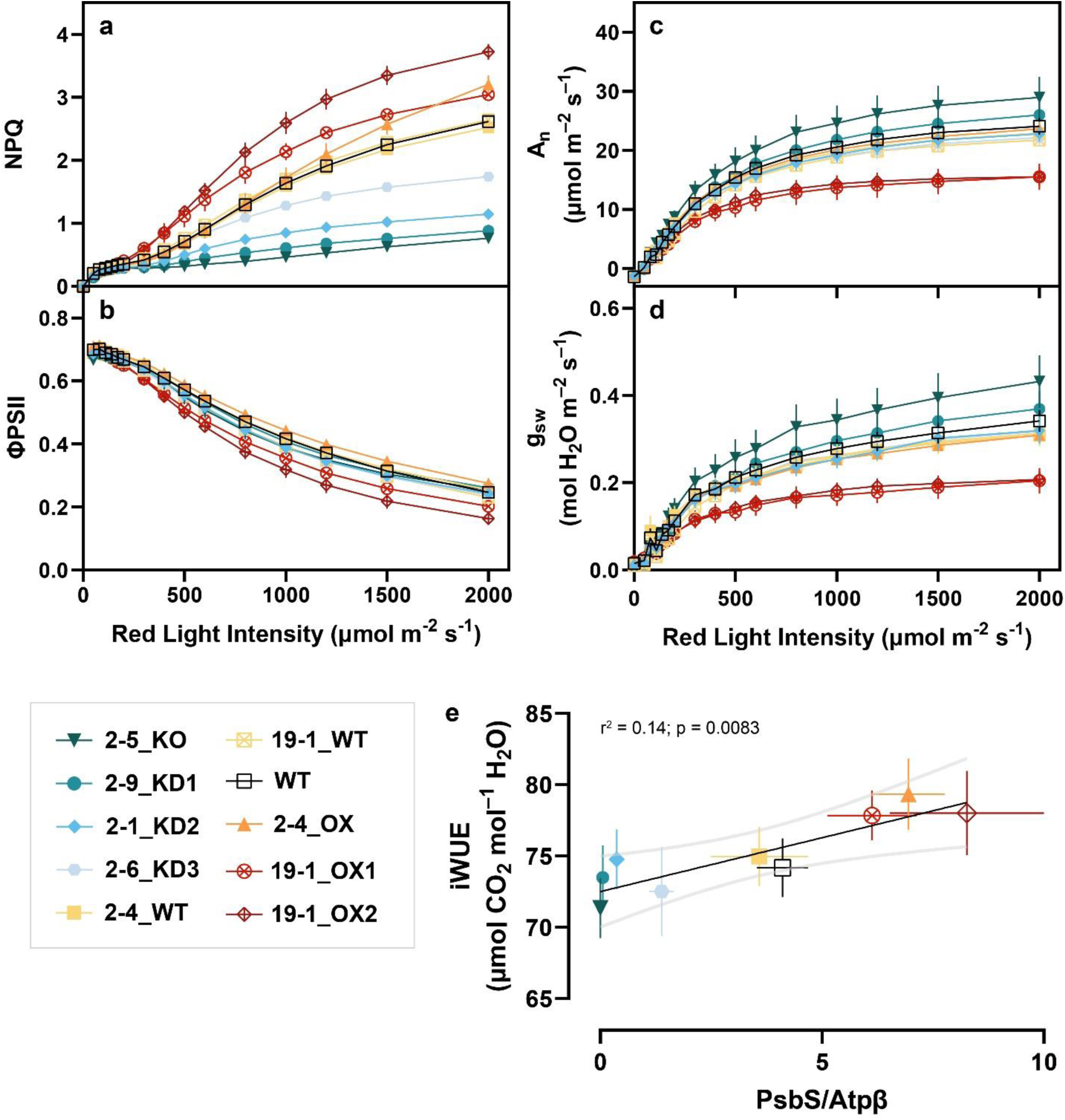
Concurrent chlorophyll fluorescence and gas exchange measurements of representative OsPSBS1-edited lines. **(a)** NPQ **(b)** operating efficiency of PSII (ΦPSII) **(c)** CO_2_ assimilation (A_n_), and **(d)** stomatal conductance (g_sw_) as a function of incident red light on mature flag leaves (n=4-7 biological replicates each, data ± SEM). **(e)** Linear regression of iWUE (A_n_/g_sw_) as a function of normalized OsPsbS1 protein abundance for each genotype (n=4-7 biological replicates, means ± SEM shown). Genotypes are shown: WT (black, open square), 2-5_KO (dark teal, inverse triangle), 2- 9_KD1 (teal, circle), 2-1_KD2 (blue, diamond), 2-6_KD3 (light blue, hexagon), 2-4_WT (yellow, square), 19-1_WT (yellow, crossed square), 2-4_OX (orange, triangle), 19-1_OX1 (red, crossed circle), 19-1_OX2 (maroon, crossed diamond).

To assess differences in iWUE due to varying PsbS activity, all iWUE values under high light (≥ 500 µmol photons m^-2^ s^-1^) were averaged per replicate and plotted against WT-normalized PsbS protein abundance. A significant, positive correlation (p=0.0083) between PsbS abundance and iWUE was observed, with a difference in average iWUE of approximately 11% between KO and OX lines (**Fig. 7e**).

### Q_A_ redox state is an inconsistent predictor of rice g_sw_

The increased iWUE phenotype observed in tobacco overexpressing *PSBS* was hypothesized to be mediated by the Q_A_ redox state, approximated by 1-qL^12^. We examined similar correlations between 1-qL and g_sw_ using our Event 2 panel, excluding possible off-target mutations affecting fitness and gas exchange phenotypes in the 19-1_OX lines (**Fig. 7, Supplemental Fig. 7**). We observed robust, statistically significant differences in 1-qL across all non-WT genotypes at light intensities above 500 µmol photons m^-2^ s^-1^ (**Fig. 8a**).

**Figure 8.**
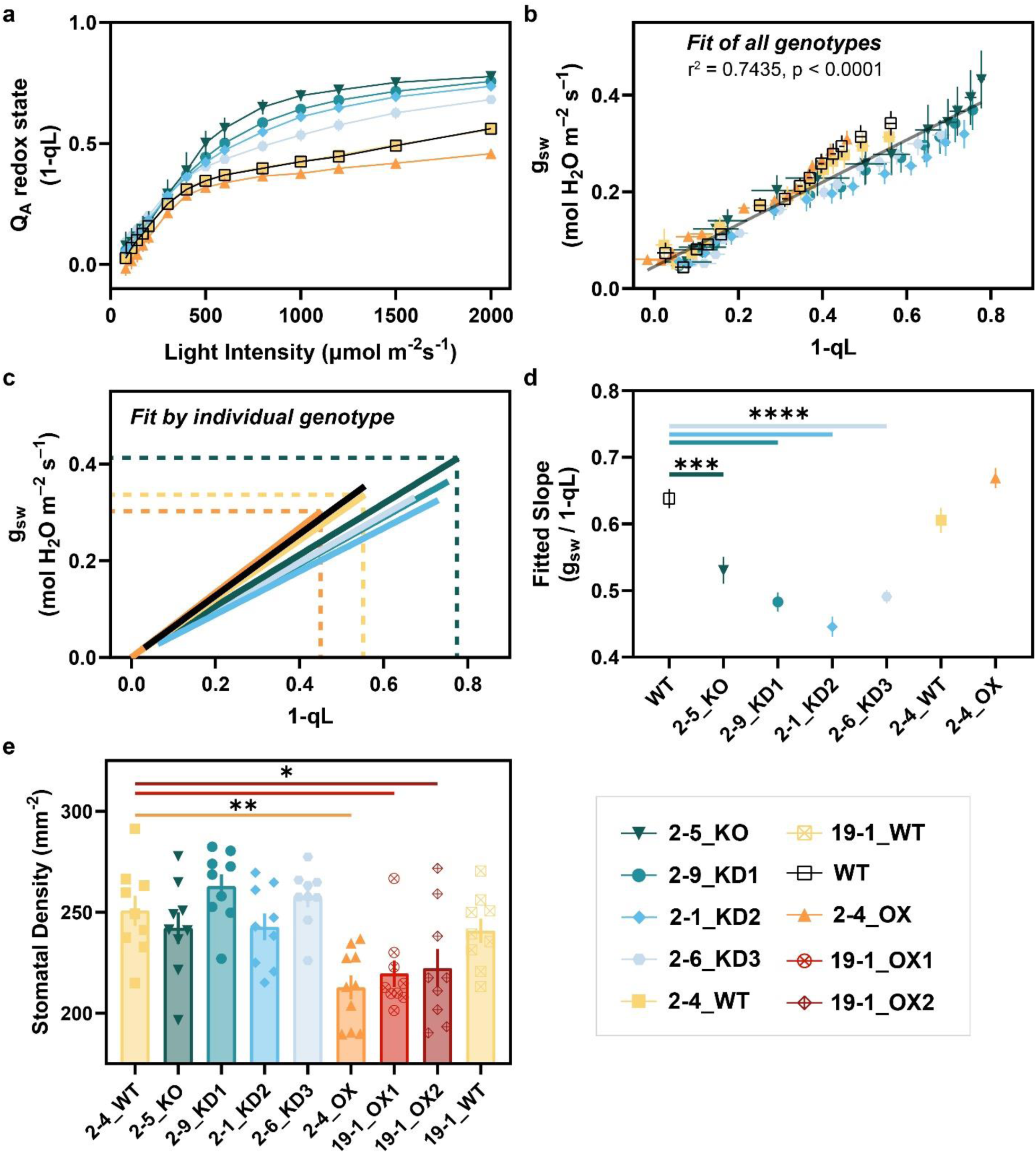
Correlation of Q_A_ redox state (1-qL) and g_sw_ as a predictor of iWUE within Event 2. **(a)** Q_A_ redox state (1-qL) as a function of incident red light on mature flag leaves (n=4-7 biological replicates, data shown ± SEM). **(b)** Linear regression of 1-qL as a function of g_sw_ for all genotypes (n=40 biological replicates) and **(c)** individuals genotypes (n=4-7 biological replicates each) at all light steps exceeding 50 µmol m^-2^ s^-1^ (data shown ± SEM). Dashed lines indicate 1-qL and gsw values at 2000 µmol photons m^-2^ s^-1^ for the 2-5_KO, 2-4_WT, and 2-4_OX genotypes. **(d)** Slopes of linear regression lines by genotype (data shown ± SEM). **(e)** Stomatal densities of all gene-edited lines (n=9 biological replicates each). Genotypes across (a-e) are shown: WT (black, open square), 2-5_KO (dark teal, inverse triangle), 2-9_KD1 (teal, circle), 2-1_KD2 (blue, diamond), 2-6_KD3 (light blue, hexagon), 2-4_WT (yellow, square), 19-1_WT (yellow, crossed square), 2-4_OX (orange, triangle), 19-1_OX1 (red, crossed circle), 19-1_OX2 (maroon, crossed diamond). Pairwise significance in (d,e) was determined by ordinary one-way ANOVA (α0.05) using Dunnett’s test for multiple comparisons against Nipponbare WT or 2-4_WT and is denoted by asterisks (*p≤0.05, **p≤0.01, ***p≤0.001, ****p<0.0001).

To interrogate the putative correlation, 1-qL was plotted against stomatal conductance for all genotypes and light intensities. A significantly non-linear (p<0.0001) and relatively strong goodness of fit (r^2^ = 0.744) was observed (**Fig. 8b**), though the correlation was weaker than what was reported in tobacco (r^2^=0.98, p<0.001)^12^. We leveraged the extended dynamic range in 1-qL phenotypes generated within this study to analyze linear regressions between 1-qL and g_sw_ by individual genotype. While higher 1-qL values were correlated with higher maximum g_sw_ values (dashed lines), the slopes of these regressions varied considerably across genotypes (**Fig. 8c**). In fact, there were statistically significant decreases in the relationship between g_sw_ and 1- qL (i.e. slope) in all KO and KD lines relative to WT. In other words, larger changes in 1-qL are associated with smaller relative changes in stomatal conductance in KO/KD but not WT/OX genotypes (**Fig. 8d**).

Finally, we measured stomatal density (SD) across all gene-edited lines. Increases in PsbS protein abundance were correlated with statistically significant decreases in stomatal density of approximately 10% (0.01 < p < 0.05) (**Fig. 8e**), with no significant differences in KO or KD lines.

## Discussion

As our understanding of basic biology grows, so does the potential for specific gene-editing solutions to agricultural and medicinal challenges. The CRISPR toolkit has the potential to meet that need if the design principles involved in the modulation of gene expression are well understood. Here we show that multiplexed CRISPR/Cas9 mutagenesis of non-coding sequences can achieve overexpression of endogenous genes and increase protein abundance to levels comparable to transgenic overexpression. Facilitated by a high-throughput screening pipeline for rapid detection of homozygous, Cas9-free progeny (**Fig. 1**) which revealed a diverse array of quantitative phenotypes (**Fig. 2**), these results have the potential to inform gene-editing strategies to optimize and fine-tune crop phenotypes.

The success of this screen was predicated on PsbS semi-dominance. However, this forward genetics approach came with inherent challenges. For example, the logarithmic relationship between PsbS abundance and NPQ (**Fig. 3e**) suggests that such a screen would enrich itself for KO/KD mutations, as modest gains in PsbS abundance will have negligible effects on overall NPQ capacity. While the two OX alleles identified in this study had significantly higher PsbS protein abundance and NPQ, it is possible that small-effect alleles are masked within the noise of WT-like biological variation (**Fig. 2**). Regardless, at a 1.67% observed frequency, it is important to recognize the constraints of random CRE mutagenesis for OX. Advances in our understanding of CRE gene expression signatures may aid in generating and resolving small-effect alleles.

Consistent with a large body of literature, we find that overexpression of PsbS increases NPQ capacity^11,12,27^. Overexpression of *OsPSBS1* in the 2-4_OX allele only increased steady-state NPQ capacity at very high light intensities (1500 & 2000 µmol photons m^-2^ s^-1^), similar to PsbS OX phenotypes in tobacco. Regardless, a strong correlation was observed between PsbS protein abundance and iWUE at light intensities as low as 500 µmol photons m^-2^ s^-1^ (**Fig. 7e**), suggesting that NPQ alone is not driving the iWUE response. Several converging lines of evidence, reviewed by Busch^28^, suggest the redox state of plastoquinone may be a well-correlated, red-light responsive regulator that might link photosynthesis and stomatal conductance through a yet unresolved signaling mechanism. Glowacka and Kromdijk et. al. provided supporting correlations for this hypothesis through the overexpression of PsbS in tobacco^12^, sidestepping pleiotropic effects observed in g_sw_ correlative studies that knocked down electron transport and CO_2_ assimilation^29,30^. However, in rice the increased WUE phenotype is poorly explained by Q_A_ redox state, as assessed by 1-qL (**Fig. 8**), which varied in its correlation with stomatal conductance across genotypes.

We also observed a marked decrease in stomatal density (SD) across all three OsPsbS1 OX lines (**Fig. 8e**), suggesting the observed differences in iWUE cannot be explained by differences in stomatal aperture alone. There is abundant historical evidence that environmental cues such as light intensity, CO_2_ availability, and water stress regulate stomatal development^31,32^. Is it possible that the overexpression of PsbS, and its downstream effects on light harvesting, intersect with some of these pathways? We cannot exclude the possibility that cumulative differences in excitation pressure at PSII during development (e.g. within vulnerable, emerging leaves), potentially signaled by Q_A_ redox state, impact iWUE through a light-dependent effect on stomatal development and aperture. However, the lack of a correlative increase in SD in any of the KO or KD lines suggests 1-qL alone is not a sufficient proxy for photosynthesis-dependent iWUE in our system (**Fig. 8e**). Morphological differences, such as the kinetically faster^33^ dumbbell-shaped stomata in monocots^34^, may also contribute to the reduced Q_A_-g_sw_ correlation and the relatively modest gains in iWUE (average ∼11% over KO, ∼7% over WT) within the already more water-use efficient grasses.

While the observed phenotypes observed reinforce expected differences in PsbS-dependent traits, the most important results of this study relate to the structural variants underlying changes in gene expression. Due to the distribution of gRNA sites in our multiplexed experimental design, we found that distal regions of the *OsPSBS1* promoter, which are conserved between *indica* and *japonica* subspecies (**Supplementary Table 1**), were dispensable for expression of *OsPSBS1*, and much of the phenotypic variation observed was due to indels near and within the 5’UTR (**Fig. 4b**). The phenotypic consequences of editing monocot CNS underscore results from other leading works that have also converged on the importance and complexity of CNS editing^19,35^.

Interestingly, sequencing of our knockdown and knockout alleles point to a possible role of a HY5-like ortholog, bZIP18, and other partially redundant bZIPs in being necessary for *OsPsbS1* expression (**Fig. 4d**). Rice bZIP18 may function similarly to Arabidopsis HY5 in partially regulating light-dependent gene expression^36^, though whether *OsPSBS1* mRNA expression is regulated by differences in TF binding and transcription rate, changes in transcript stability, or other limiting mechanisms remains to be determined. Regardless, the reduced genetic complexity (e.g. need for fewer guides, smaller editing region) presented by 5’UTR mutagenesis of sites flanking conserved non-coding sequences may be an attractive approach for achieving controlled knockdown to bypass negative epistasis^37^. The potential of 5’UTR mutagenesis for obtaining overexpression alleles, including the editing of competitive upstream open reading frames (ORFs) that may inhibit translation^38^ or the use of prime editing approaches^39^ to introduce small enhancing GATC-motifs downstream of the TSS^40^, warrants further investigation.

Surprisingly, the two overexpression alleles were caused by inversions upstream of *OsPsbS1* (**Fig. 5**). Allele 19-1_OX1 presents an interesting example as it carries a clean inversion between a pair of distal and proximal gRNAs that may readily be replicated or introgressed into other varieties. Unfortunately, the persistence of the high copy number T-DNA transgene in this line may contribute to the decreased overall fitness of 19-1 OX progeny (**Supplementary Fig. 7**). These pleiotropic off-target effects are also reflected in reductions in 19-1_OX gas exchange and ΦPSII phenotypes (**Fig. 7b-d**), and the significantly increased NPQ capacity at lower light intensities (**Fig. 7a**), all of which have not been reported in transgenic rice PsbS overexpression experiments^11^ or seen in the 2-4_OX line that displays a similar magnitude of PsbS1 overexpression (**Fig. 3d**). Future work to isolate this allele in a Cas9-free background, or recapitulate the allele in other varieties of rice, will be worthwhile to de-convolute PsbS abundance, photosynthesis, and gas exchange phenotypes.

The 2-4_OX 252-kb duplication/inversion was an unexpected, but interesting result from this screen for transgene-free overexpression. Despite this significant genomic perturbation, we observed negligible changes in global gene expression (**Fig. 6**) with most of the DEG within the duplication/inversion itself. Correlating with copy number, we observe 2-fold overexpression of not just *OsPSBS1*, but also the remaining genes within the duplication. The OX of the remaining 35 genes may result in other gain-of-function phenotypes that were not resolved within our NPQ-focused screen. KEGG enrichment of photosynthesis and carbon fixation metabolic pathways suggests that overexpression of PsbS, and potentially its adjacent genes, may have synergistic effects in contributing towards overall photosynthetic efficiency.

A revolution in long-read sequencing has revealed the pervasiveness of genomic structural variants. Complex structural variants (CSVs) including translocations, insertions, and inversions are persistent at both the population and pan-genome level as reported in tomato^41^, rapeseed^42^, maize^43^, grapevine^44^, and rice^45^, affecting upwards of 20% of all genes. In fact, much of the current understanding of structural variants is driven by the prevalence of these changes driving various human cancers and diseases^46–48^. Unlike humans however, plants exhibit a much greater tolerance to (and thus abundance of) structural variants that is likely driven *in vivo* by transposable elements and recombination—both documented as key drivers of crop domestication^49,50^.

In fact, chromosomal inversions have also been implicated in gene overexpression during the domestication of peach^51^ and the complex rearrangements that underlie multiple myeloma^52^. Recently, Lu et. al. showed that CRISPR/Cas9 could be used to drive native overexpression via promoter swapping^20^, generating inversions in ∼3% of transformed calli that increased gene expression of *OsPPO1* at varying frequencies. However, these inversions came at the cost of expression of the opposite promoter, knocking out expression of the Calvin-Benson cycle protein 12 (*OsCP12)* gene (LOC_Os01g19740). In the case of our 19-1_OX1 allele, the other inversion breakpoint occurred downstream of the 3’UTR of the neighboring gene (LOC_Os01g64930) and likely does not interfere with its expression. The work presented here, unbiased in its target design, reinforces and expands the genome-engineering potential of inversions for native gene overexpression.

In our dataset, overexpression alleles account for 2 of 13 unique alleles that could not be amplified by PCR and sequenced. Of the remaining 11 alleles, 8 were phenotypic knockouts, 1 was a phenotypic knockdown, and 2 were WT-like in NPQ activity. It is possible that complex structural variants also underlie these remaining alleles, increasing potential CSV frequency from 1.7% to 10.8%. We see significant potential in expanding beyond TILLING (Targeting Induced Local Lesions in Genomes) and local CRE-editing Cas9-mediated approaches to drive significant rather than small-effect changes in gene expression phenotype. Efforts to increase the efficiency and frequency of CSVs such as by use of alternate Cas systems^19^, paired prime-editing strategies^53^ or DNA repair mutants backgrounds^54^ may be enticing options to enrich for significant genomic disruptions and changes in endogenous gene overexpression. Guided design of inversions to minimize transcriptomic perturbations, such as inversions into transcriptionally active epigenetic marks^55^ rather than promoter swaps, may also prove fruitful in future plant engineering efforts. Additionally, investigating the potential of editing other non-coding sequence types (e.g. introns and 3’UTRs) may expand gene-editing opportunities for fine-tuning expression. Further research identifying promising genomic markers for putative overexpression, such as synthetic enhancer signatures^56^ and high expression 5’UTR motifs^40^, are necessary to expand the reproducibility of these efforts. Applying these hypotheses to other NPQ genes, such as *OsVDE* and *OsZEP*, will bring us closer to optimizing photoprotection and native photosynthetic efficiency in crop plants.

## Materials and Methods

### Plant material and growth conditions

Rice cultivar Nipponbare (*Oryza sativa sp. japonica*) seeds were germinated on Whatman filter paper for five days at 100 μmol photons m^-2^ s^-1^ fluorescent light with a 14-hour day length (27°C day/25°C night temperature). Seedlings were transferred to soil composed of equal parts Turface and Sunshine Mix #4 (Sungro) and grown under seasonal day length (10-14 hours) in a south-facing greenhouse that fluctuated in temperature (38°C High/16°C Low) and relative humidity (45-60%). Plants were fertilized with a 0.1% Sprint 330 iron supplement after transplanting at 2 weeks post-germination and at the onset of grain filling at 10 weeks post germination, and JR Peter’s Blue 20-20-20 fertilizer biweekly. Flats were kept full of water to mimic flooded growth conditions. Genotypes were randomized across flats and throughout the greenhouse to minimize positional effects. At the V4-5 leaf stage, T_1_ progeny and WT controls were assayed for differences in NPQ capacity and sensitivity to the selectable marker hygromycin.

To assess differences in photosynthetic efficiency and yield, gene-edited T_3_ plants were grown in a larger greenhouse with more homogenous light exposure and field-relevant growth conditions (40°C High/27°C Low, 30-50% relative humidity) without supplemental light.

### Guide RNA design and cloning into Cas9 vector pRGEB32

Eight gRNA target sites were identified upstream of the functional *PsbS* ortholog in rice, *PsbS1* (LOC_Os01g64960), using CRISPR-P (<u>crispr.hzau.edu.cn</u>)^57^ and a 1.5-kb region upstream of the *PsbS1* start codon in a draft genome of *Oryza sativa sp. indica* cultivar IR64 (http://schatzlab.cshl.edu/data/rice/)^58^ (**Supplementary Table 1**). The eight gRNA spacers were assembled into a DNA cassette interspersed with scaffolds and tRNA linkers for polycistronic gRNA expression as previously described^59^, and synthesized (Genscript). The insert was cloned into the pRGEB32 rice *Agrobacterium*-mediated transformation vector (Addgene Plasmid #63142) via GoldenGate Assembly.

### Induction of embryogenic calli

Mature seeds of rice (*Oryza sativa ssp. japonica* cv. Nipponbare) were de-hulled, and surface-sterilized for 20 min in 20% (v/v) commercial bleach (5.25% sodium hypochlorite) plus a drop of Tween 20. Three washes in sterile water were used to remove residual bleach from seeds. De-hulled seeds were placed on callus induction medium (CIM) medium (N6 salts and vitamins^60^, 30 g/L maltose, 0.1 g/L myo-inositol, 0.3 g/L casein enzymatic hydrolysate, 0.5 g/L L-proline, 0.5 g/L L-glutamine, 2.5 mg/L 2,4-D, 0.2 mg/L BAP, 5 mM CuSO_4_, 3.5 g/L Phytagel, pH 5.8) and incubated in the dark at 28 °C to initiate callus induction. Six-to eight-week-old embryogenic calli were used as targets for transformation.

### Agrobacterium-mediated transformation

Embryogenic calli were dried for 30 min prior to incubation with an *Agrobacterium tumefaciens* EHA105 suspension (OD_600nm_ = 0.1) carrying the cloned binary vector, pRGEB32_OsPsbS1_8xgRNA. After a 30 min incubation, the *Agrobacterium* suspension was removed. Calli were then placed on sterile filter paper, transferred to co-cultivation medium (N6 salts and vitamins, 30 g/L maltose, 10 g/L glucose, 0.1 g/L myo-inositol, 0.3 g/L casein enzymatic hydrolysate, 0.5 g/L L-proline, 0.5 g/L L-glutamine, 2 mg/L 2,4-D, 0.5 mg/L thiamine, 100 mM acetosyringone, 3.5 g/L Phytagel, pH 5.2) and incubated in the dark at 21°C for 3 days. After co-cultivation, calli were transferred to resting medium (N6 salts and vitamins, 30 g/L maltose, 0.1 g/L myo-inositol, 0.3 g/L casein enzymatic hydrolysate, 0.5 g/L L-proline, 0.5 g/L L-glutamine, 2 mg/L 2,4-D, 0.5 mg/L thiamine, 100 mg/L timentin, 3.5 g/L Phytagel, pH 5.8) and incubated in the dark at 28°C for 7 days. Calli were then transferred to selection medium (CIM plus 250 mg/L cefotaxime and 50 mg/L hygromycin B) and allowed to proliferate in the dark at 28°C for 14 days. Well-proliferating tissues were transferred to CIM containing 75 mg/l hygromycin B. The remaining tissues were subcultured at 3- to 4- week intervals on fresh selection medium. When a sufficient amount (about 1.5 cm in diameter) of the putatively transformed tissues was obtained, they were transferred to regeneration medium (MS salts and vitamins ^61^, 30 g/L sucrose, 30 g/L sorbitol, 0.5 mg/L NAA, 1 mg/L BAP, 150 mg/L cefotaxime) containing 40 mg/L hygromycin B and incubated at 26 °C, 16-hr light, 90 µmol photons m^-2^ s^-1^. When regenerated plantlets reached at least 1 cm in height, they were transferred to rooting medium (MS salts and vitamins, 20 g/L sucrose, 1 g/L myo-inositol, 150 mg/L cefotaxime) containing 20 mg/L hygromycin B and incubated at 26 °C under conditions of 16-hr light (150 µmol photons m^-2^ s^-1^) and 8-h dark until roots were established and leaves touched the Phytatray lid. When possible, multiple regenerants per calli were recovered. Plantlets were then transferred to soil.

### Chlorophyll fluorescence measurements of NPQ

Leaf punches were sampled from mature, fully developed leaves at leaf stage V3-5 and floated on 270 µL of water in a 96-well plate. Plates were dark acclimated for at least 30 minutes prior to analysis. *In vivo* chlorophyll fluorescence measurements were determined at room temperature using an Imaging-PAM Maxi (Walz) pulse-amplitude modulation fluorometer. Fluorescence levels after dark acclimation (F_o_, F_m_) and during light acclimation (F_o_’, F_m_’) were monitored in two ways:

To resolve phenotypically segregating *OsPsbS1* NCS gene-edited alleles from a single parent, a single leaf punch at the V3 stage was exposed to a 4-minute period of high-intensity actinic light (1500 µmol photons m^-2^ s^-1^) using periodic saturated pulses. Putative homozygous lines were identified as those progenies in the top and bottom 10% of total NPQ. The leaf punches were then used to determine hygromycin sensitivity and Cas9 transgene segregation as described (0).

Candidates above lacking Cas9 when possible were re-sampled at the V5 leaf stage, phenotyping two leaf punches from two mature leaves per plant to compare NPQ relative to Nipponbare WT plants. NPQ was quantified during a 10-minute period of high-intensity actinic light (1500 µmol photons m^-2^ s^-1^) and 10 minutes dark relaxation (0 µmol photons m^-2^ s^-1^) using periodic saturating pulses. NPQ in both cases was calculated (Equation 1).

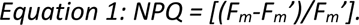

### Hygromycin sensitivity assay for high-throughput Cas9 transgene detection

Floated leaf punches assayed for NPQ capacity were used to determine hygromycin sensitivity and segregation of the transgene. Hygromycin B (50 mg/mL 1x PBS) was added to each well to a final concentration of 20 µg/mL of antibiotic. Plates were incubated under rice germination conditions for three days, following which the Imaging-PAM Maxi (Walz) was used to identify differences in the maximum efficiency of Photosystem II (F_v_/F_m_) after 30 minutes dark acclimation using (Equation 2).

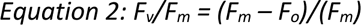

Sensitive, transgene-free plants had a decline in F_v_/F_m_ of >0.3-0.4, whereas transgenic plants maintained a WT F_v_/F_m_ of ∼0.7-0.8.

### Total RNA/protein extraction of representative lines with varying NPQ

2 cm of leaf tissue from the youngest fully developed leaf was collected and flash-frozen in RNAse-free, DNAse-free tubes containing Lysing Matrix D (FastPrep-24^TM^) at midday. Leaf tissue was ground on dry ice using a FastPrep-24 5G™High-Speed Homogenizer (6.0 m/s for 2 x 40 s, MP Biomedical). Protein and mRNA were extracted from the same leaf sample (NucleoSpin RNA/Protein kit, REF740933, Macherey-Nagel GmbH & Co., Düren, Germany).

### Quantitative RT-PCR of OsPsbS1 relative to two reference genes

Extracted mRNA was treated with DNase (ThermoFisher Scientific) and transcribed to cDNA using Omniscript Reverse Transcriptase (Qiagen) and a 1:1 mixture of random hexamers and oligo dT as recommended by the manufacturer. Quantitative reverse transcription PCR was used to quantify *OsPsbS1* transcripts relative to *OsUBQ* and *OsUBQ5* transcripts, in biological triplicate with technical duplicates using published methods to normalize qRT-PCR expression to multiple reference genes^62,63^. Samples were run on a 7500 Fast Real-Time PCR system (Applied Biosystems, Gent, Belgium) in a total volume of 20 µL using 4 µL of 1:10 diluted cDNA. Primers were empirically validated on a 5-step dilution series of WT cDNA (1:1 to 1:81). All final primer pairs had an amplification efficiency between 90-105% and linear amplification within the dynamic range tested. A single peak in melt-curve analysis was observed for each gene of interest, verifying specificity of the amplicon. The primer sets were similarly efficient and specific for *OsPSBS1* copy number experiments of RNA-free genomic DNA. Primers and primer efficiencies are shown (**Supplementary Table 2**).

### Immunoblotting of whole leaf protein extracts

Precipitated protein was resuspended in the supplied protein solubilization buffer (PSB-TCEP) and quantified using a TCA colorimetric assay^64^. Samples containing 6 µg total protein were resolved using pre-cast SDS-PAGE Any KD™ gels (BIO-RAD), transferred to a polyvinylidene difluoride membrane (Immobilon-FL 0.45 µm, Millipore) via wet transfer, and blocked with 3% nonfat dry milk for immunodetection. Membranes were cut and incubated with the following antibodies. A rabbit polyclonal antibody raised against sorghum PsbS (α- SbPsbS,DEEVTGLDKAVIQPGKGFRGALGLSE-Cys) was produced by Pacific Immunology and generously shared by Steven J. Burgess (University of Illinois) and used at a 1:2,500 dilution. A rabbit polyclonal antibody raised against a synthetic peptide of the β-subunit of ATP synthase (Atpβ) was obtained from Agrisera (catalogue no. AS05 085) and used at 1:10,000 dilution. After incubation with an HRP-conjugated, anti-rabbit secondary antibody from GE Healthcare (1:10,000 dilution), bands were detected by chemiluminescence using SuperSignal West Femto Maximum Sensitivity Substrate (Thermo Scientific). Protein bands were quantified by densitometry with ImageQuant TL software (version 7.0 GE Healthcare Life Sciences, Pittsburgh, PA, USA). PsbS abundance was quantified relative to WT based on a dilution series of WT PsbS protein.

### PCR genotyping of transgene-free, edited lines

50 mg of leaf tissue was ground via bead beating (Lysing Matrix D, FastPrep-24^TM^) and genomic DNA was extracted in 2xCTAB buffer at 65°C for 15 minutes. DNA was separated via chloroform phase separation and precipitated using isopropanol. The pellet was washed briefly in 70% ethanol before being dried and resuspended in 1xTE buffer.

Two overlapping regions spanning the 8 gRNA target sites were PCR-amplified via Phusion^TM^ High-Fidelity PCR using 5x GC-rich buffer (**Supplementary Table 3**). PCR products were amplified from at least two putative homozygotes per line and sequenced by Sanger Sequencing. Contigs of the ∼4.3-kb upstream region were assembled using Snapgene and analyzed by multiple sequence alignment.

### In-parallel gas exchange and chlorophyll fluorescence analysis

Photosynthetic gas exchange dynamics were measured on the youngest, fully expanded flag leaf of 12-week-old flowering rice plants. Gas exchange measurements were performed using an open gas exchange system (LI6800, LI-COR, Lincoln, NE, USA) equipped with a 2-cm^2^ leaf chamber and integrated modulated fluorometer. Whole plants were low light acclimated for 1- 2 hours to mitigate afternoon depression of photosynthesis and dark acclimated for at least 30 minutes to allow for concurrent phenotyping of gas exchange (e.g. A_n_, g_sw_) and chlorophyll fluorescence parameters (e.g. F_v_/F_m_, NPQ). For all measurements the chamber conditions were set to: 400 ppm chamber [CO_2_], 27°C chamber temperature, 1.3-1.5 kPa vapor pressure deficit of the leaf, 500 µmol s^-1^ flow rate, and a fan speed of 10,000 rpm. Samples were assayed within the boundaries of ambient daylength (8 am – 5 pm).

Steady state photosynthesis and stomatal conductance was monitored in response to changes in red light intensity (100% red LED’s, λ_peak_ 630 nm). Light intensity was varied from 0, 50, 80, 110, 140, 170, 200, 300, 400, 600, 800, 1000, 1200, 1500, and 2000 µmol photons m^-2^ s^-1^ with 10-20 minutes of acclimation per light step. Steady state was reached when the stomatal conductance, g_sw_, maintained a slope less than 0.005 +/- 0.00025 SD over a 40 second period and when net assilimation rate, A_n_, showed variation less than 0.5 +/- 0.25 SD over a 20 s period. Net assimilation rate, stomatal conductance, and intracellular [CO2] was logged. A saturating pulse was then applied to collect all relevant chlorophyll fluorescence parameters using a multiphase flash routine. In addition to NPQ and F_v_/F_m_, we assessed ΦPSII, intrinsic water use efficiency (iWUE), and Q_A_ redox state (1-qL) (Equations 3-5). The derivation of 1-qL assumes a “lake” model for photosynthetic antenna complexes.

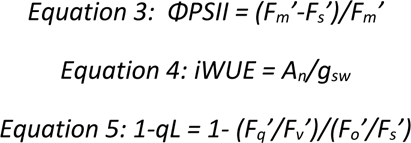

### Pacbio long-read whole-genome sequencing and analysis

Leaf tissue used for high-molecular weight genomic DNA (HMW gDNA) extraction was dark starved for 4 days before being sampled and flash frozen in liquid nitrogen. DNA was isolated using the NucleoBond HMW DNA kit (TakaraBio, Catalog #740160.2) with the following modifications: Plant leaves were ground by pestle and mortar under liquid nitrogen, where 1 g of ground leaf tissue was resuspended in 2.5 times the amount of recommended lysis buffer and incubated in a 50°C water bath for 4 hours. The amount of Binding Buffer H2 was also proportionately increased. HMW gDNA was resuspended by gentle pipetting in water and assessed for quality by Femto Pulse Analysis (median fragment length 13-15 kb).

HMW gDNA was pooled and sequenced using a PacBio Sequel II (QB3 Genomics, UC Berkeley, Berkeley, CA, RRID:SCR_022170) by HiFi circular consensus sequencing (CCS) on a single 8M SMRTcell. Untrimmed reads for overexpression lines N2-4_OX and N19-1_OX were first quality checked using FastQC (https://www.bioinformatics.babraham.ac.uk/projects/fastqc/). Reads were mapped to the *Oryza sativa* v7.0 reference genome^24^ using pbmm2, a minimap2 wrapper by Pacific Biosciences specifically for HiFi reads and assembled using pbIPA, Pacific Biosciences phased assembler (https://github.com/PacificBiosciences/pbbioconda). Sniffles (https://github.com/fritzsedlazeck/Sniffles) and pbsv (https://github.com/PacificBiosciences/pbbioconda) were used to identify structural variants from mapped reads. D-Genies^65^ was used to align the de novo assembly to the reference genome and generate dot plots. Integrative Genomics Viewer^25^ was used to visualize mapped reads, aligned assemblies, and sniffles structural variation output.

### Whole transcriptome sequencing

RNAseq was performed by Novogene (www.novogene.com). Briefly, messenger RNA was purified from total RNA using poly-T oligo-attached magnetic beads for 150 bp paired-end sequencing on a NovaSeq6000. Raw data (raw reads) in fastq format was first processed using Novogene perl scripts. Reference genome and gene model annotation files were downloaded from Phytozome^24^. Paired-end clean reads were aligned to an index of the reference genome built using Hisat2 v2.0.5^66^. The mapped reads of each sample were assembled by StringTie v1.3.3b^67^ in a reference-based approach. FeatureCountsv1.5.0-p3^68^ was used to count the reads numbers mapped to each gene. Then FPKM of each gene was calculated based on the length of the gene and reads count mapped to this gene. Fold change differences were calculated using the DESeq2 package^69,70^. The resulting p-values were adjusted using the Benjamini and Hochberg’s approach for controlling the false discovery rate. Genes with an adjusted p-value <=0.10 found by DESeq2 were assigned as differentially expressed. Finally, we used the clusterProfiler R package to test the statistical enrichment of differential expression genes in KEGG pathways^71^.

### Stomatal density measurements

Stomatal density was measured on the abaxial side of the fourth fully expanded true leaf with method adapted from Karavolias et al. 2023^72^. Briefly, epidermal impressions of nine biological replicates were taken from the widest section of the leaves. Images of each impression were taken using a Leica DM5000 B epifluorescent microscope at 10x magnification. Three images were captured from each impression. The number of stomatal in a single stomatal band was counted and divided by the area of the band to calculate stomatal density. The stomatal density calculated from the three images was averaged to represent the stomatal density of each biological replicate.

## Acknowledgements

We thank Steven Burgess and Lynn Doran (University of Illinois) for generously providing the Sorghum PsbS antibody used in this study. We thank Christina Wistrom, Bradlee McAnally, and the Oxford Tract greenhouse staff for their assistance in plant maintenance. This work was supported by a sub-award from the University of Illinois as part of the research project Realizing Increased Photosynthetic Efficiency (RIPE) that is funded by the Bill & Melinda Gates Foundation, Foundation for Food and Agriculture Research, and the U.K. Foreign, Commonwealth & Development Office under grant number OPP1172157. D.P-T. was supported by the Berkeley Fellowship and the NSF Graduate Research Fellowship Program (Grant DGE 1752814). K.K.N. is an investigator of the Howard Hughes Medical Institute. This article is subject to HHMI’s Open Access to Publications policy. HHMI lab heads have previously granted a nonexclusive CC BY 4.0 license to the public and a sublicensable license to HHMI in their research articles. Pursuant to those licenses, the author-accepted manuscript of this article can be made freely available under a CC BY 4.0 license immediately upon publication.

## Author Contributions

D.P-T. and K.K.N. designed experiments. D.P-T. designed the editing plasmid. M.T. and M-J.C. generated transgenic rice lines. D.P-T. designed the NPQ and Cas9 screen. A.K. screened the rice mutant library and managed all plants. D.P-T. and A.L. genotyped mutants and quantified changes in gene expression. D.P-T. quantified changes in protein abundance. A.K. and A.L. measured dried biomass. N.M. analyzed the 5’UTR sequencing and long-read sequencing data. D.P-T. analyzed RNAseq data generated under contract with Novogene (www.novogene.com). D.P-T. and A.K. performed gas exchange measurements. N.G.K. performed stomatal density measurements. D.P-T. analyzed experiments and wrote the manuscript with input from K.K.N.

