## Supplemental Data for "Multiplexed CRISPR/Cas9 mutagenesis of rice PSBS1 non-coding sequences for transgene-free overexpression"

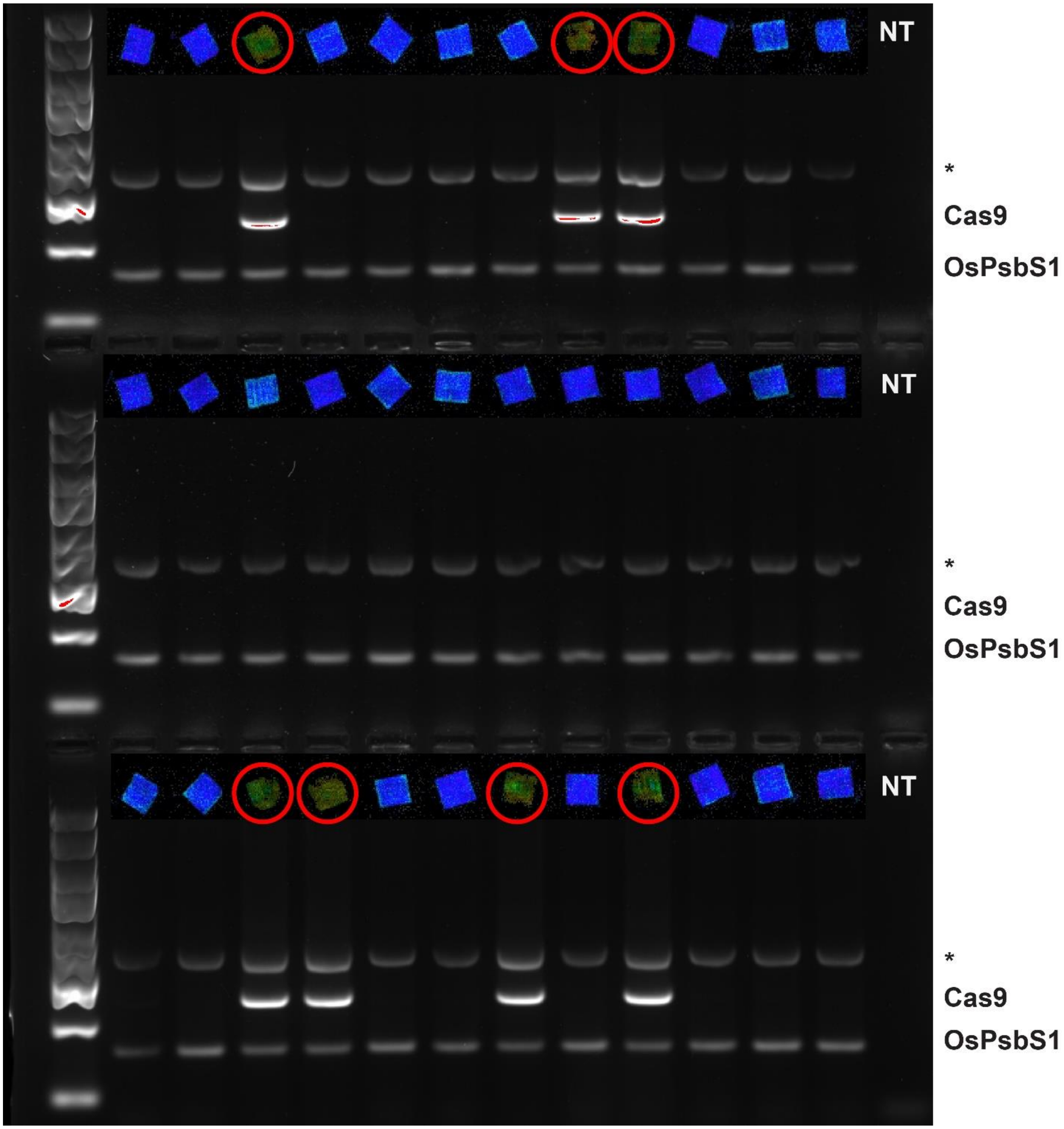


***Supplementary Figure 1.*** *Transgene genotyping of hygromycin-resistant and -sensitive plants.*

Gel electrophoresis results of pooled PCRs amplifying the *Cas9* transgene and a region within the *OsPsbS1* coding sequence. Corresponding leaf punches from (Fig. 1f) are shown above each lane, with leaves with reduced $F_v/F_m$ that are sensitive to the antibiotic are circled in red. A non-specific background band is noted by an asterisk (*).

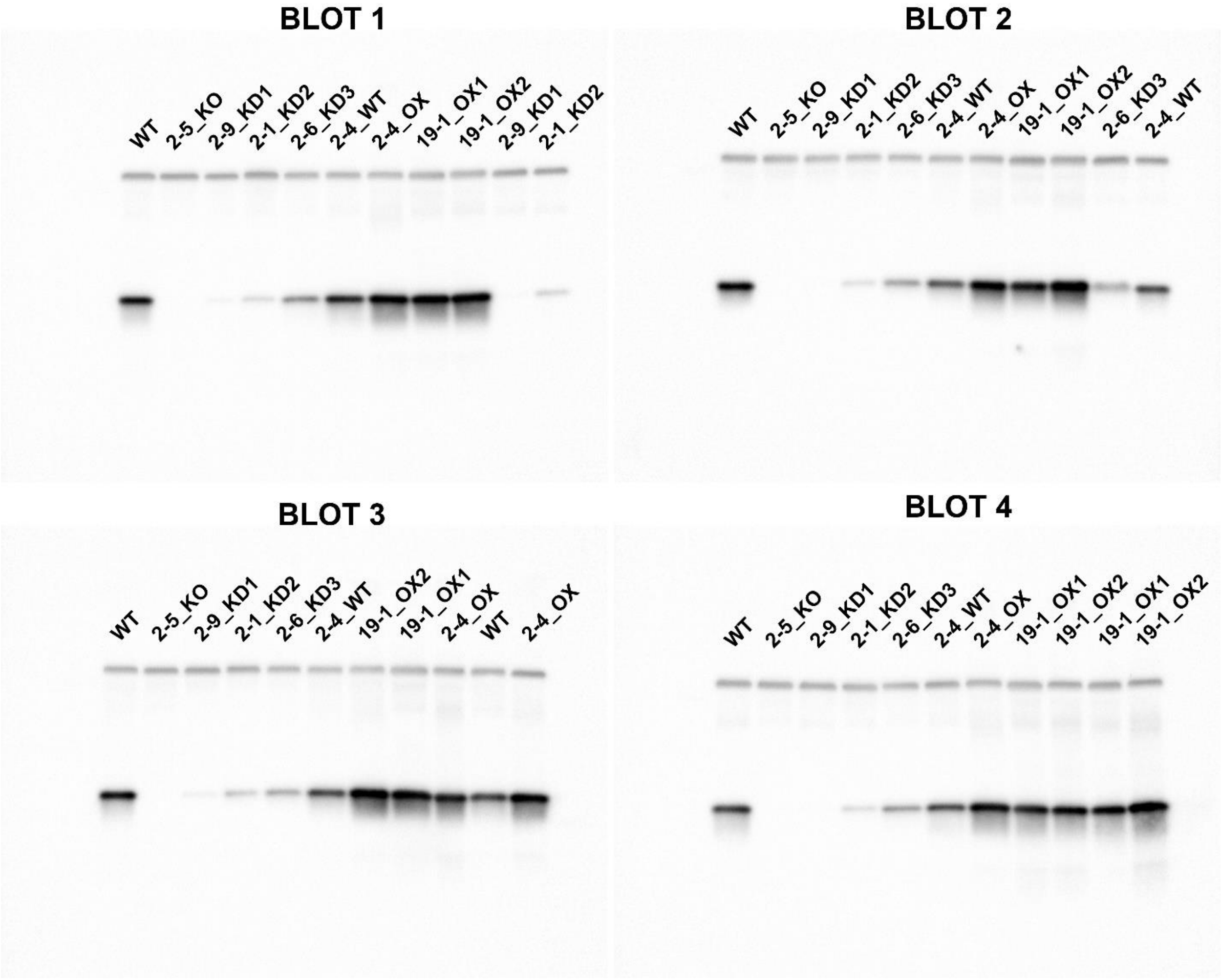


***Supplementary Figure 2.*** *All immunoblots used for quantification of Atpβ and PsbS.*

Immunoblots detecting AtpB (1:10000, top band) and PsbS (1:2500, bottom band) on 4 independent blots spanning 4-5 biological replicates per genotype at a 5 s exposure time. 6 µg of total protein was loaded into each well.

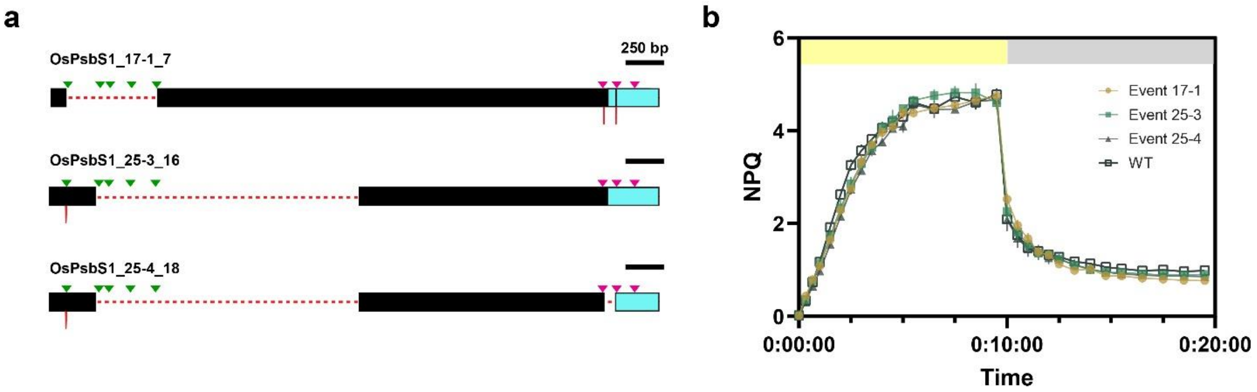


***Supplementary Figure 3.*** *Large distal deletion alleles and associated NPQ capacity.*

**(a)** Three unique cis-regulatory mutants with large deletions (red dashed lines) at distal gRNA sites (green triangles) mapped onto the Nipponbare promoter. 5'UTR is shown in cyan. Scale bar is 250 bp. **(b)** NPQ kinetics of large distal deletion lines in (a) as follows: Event17-1_7 (brown, circle), Event 25-3_16 (teal, square), Event 25-4_18 (gray, triangle), Nipponbare WT (black, open square). Data shown ± SEM.

**a**

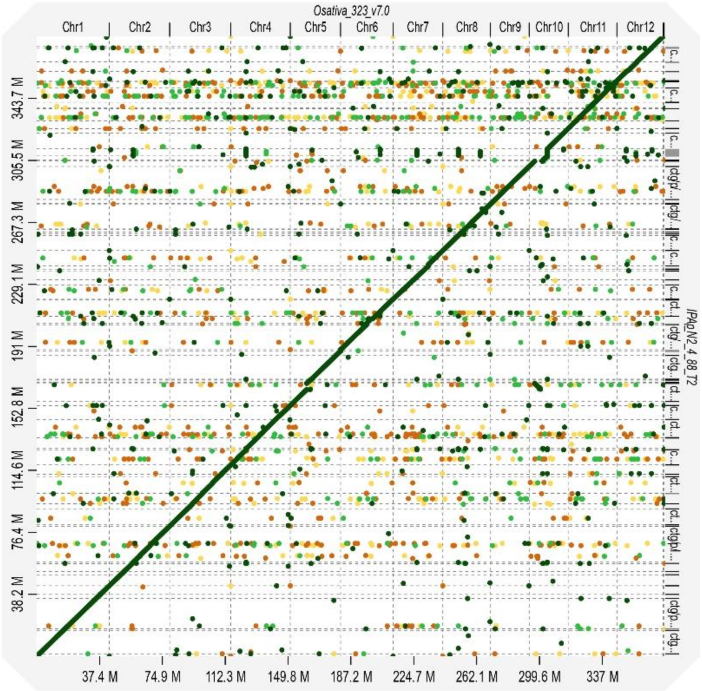


**b**

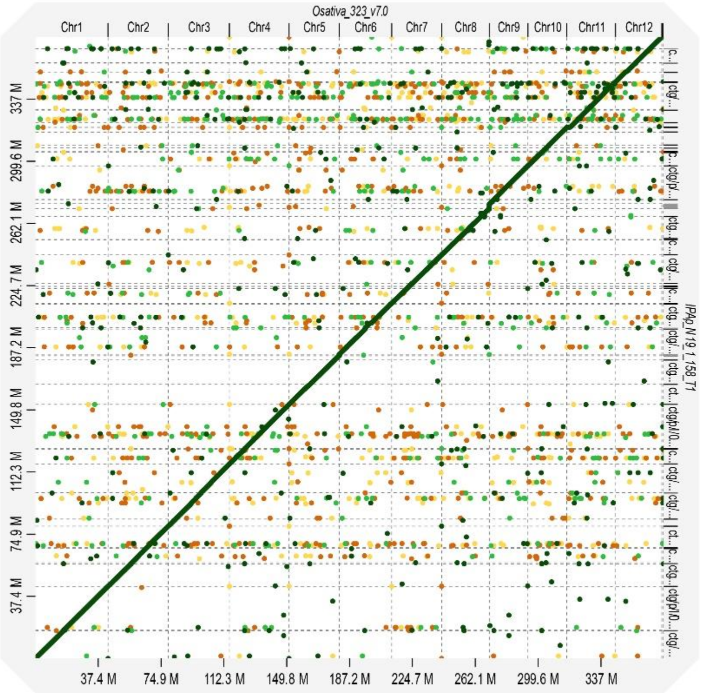


***Supplementary Figure 4****. Dot plots of Pacbio long-read overexpression allele sequencing.*

**(a)** Dot plot of 2-4_OX and **(b)** 19-1_OX, plotting sequenced OX variants (y-axis) against the reference genome (x-axis). Colors indicate the strength of matched sequences, with yellow being the lowest confidence (0-25%) and dark green being the highest (75-100%). Continuity on the intersecting axis (diagonal green) indicates high similarity between both genomes, with chromosome-level insertions and deletions shown by gaps in continuity.

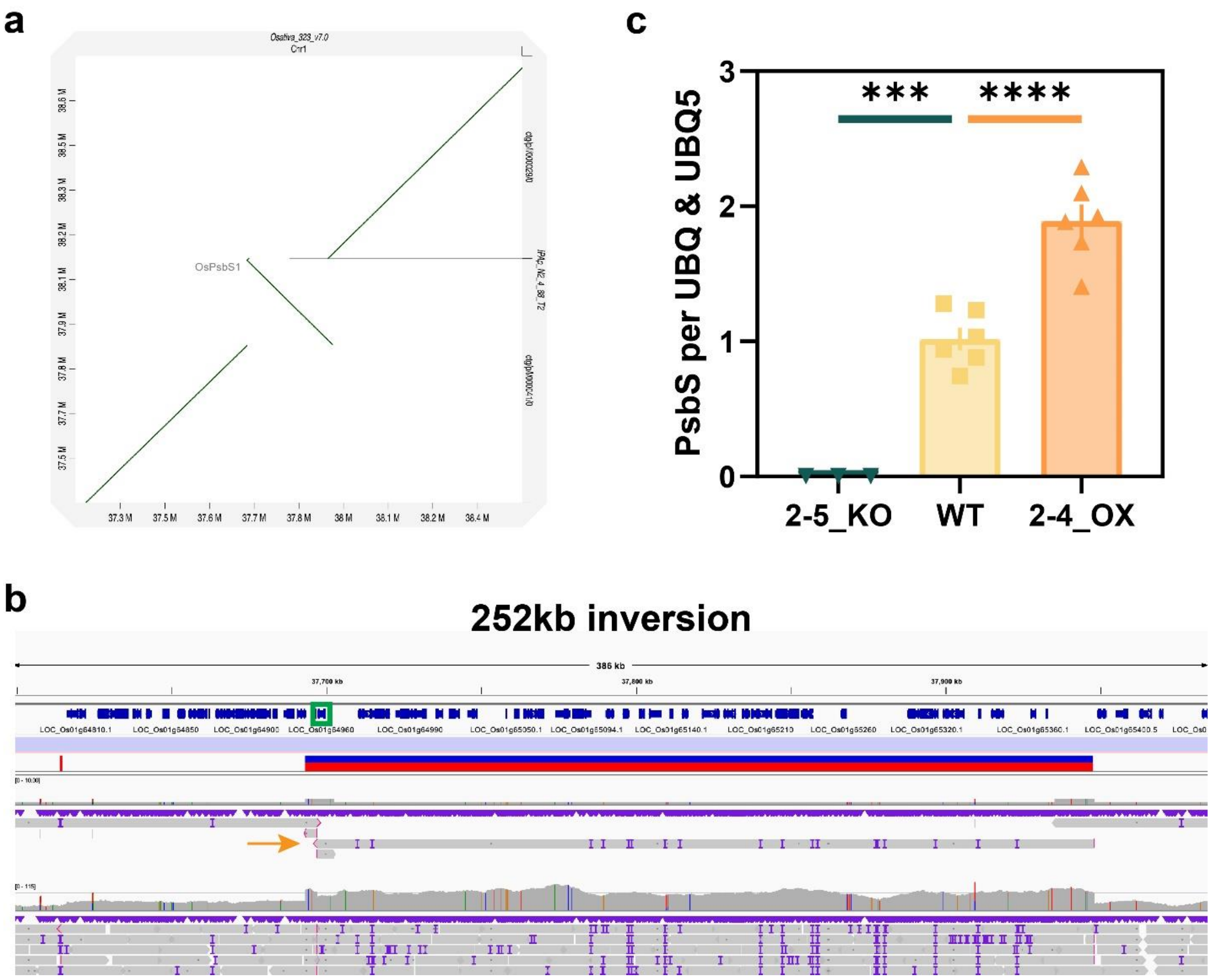


***Supplementary Figure 5.*** *High resolution view of the 2-4_OX duplication/inversion.*

**(a)** Increased resolution of the inversion on the Chr 1 locus (~37.85 Mbp). **(b)** Integrative Genomics Viewer (IGV) display of raw mapped reads. The green box denotes the *OsPSBS1* gene (LOC_Os01g64960). Structural variant called by Sniffles is shown by the red and blue bar. Evidence of the inversion is substantiated by head-to-head sequenced fragments (bottommost orange arrow). **(c)** Quantitative PCR analysis to determine PsbS copy number relative to WT and the 2-5 *OsPSBS1* deletion line. Pairwise significance in (c) was determined by ordinary one-way ANOVA ($\alpha$=0.05) using Dunnett's test for multiple comparisons against Nipponbare WT and is denoted by asterisks (***$p \leq 0.001$, ****$p < 0.0001$).

**a**

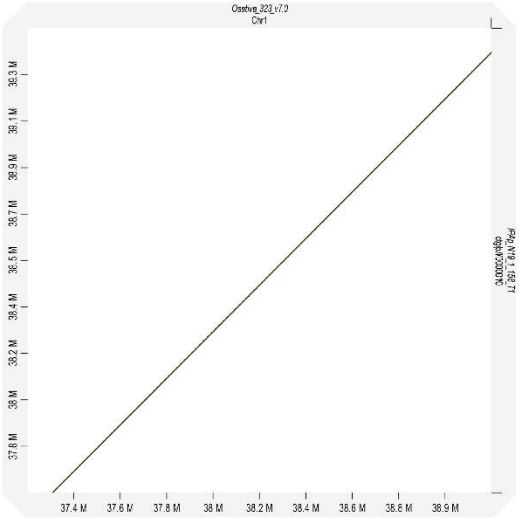

**b**

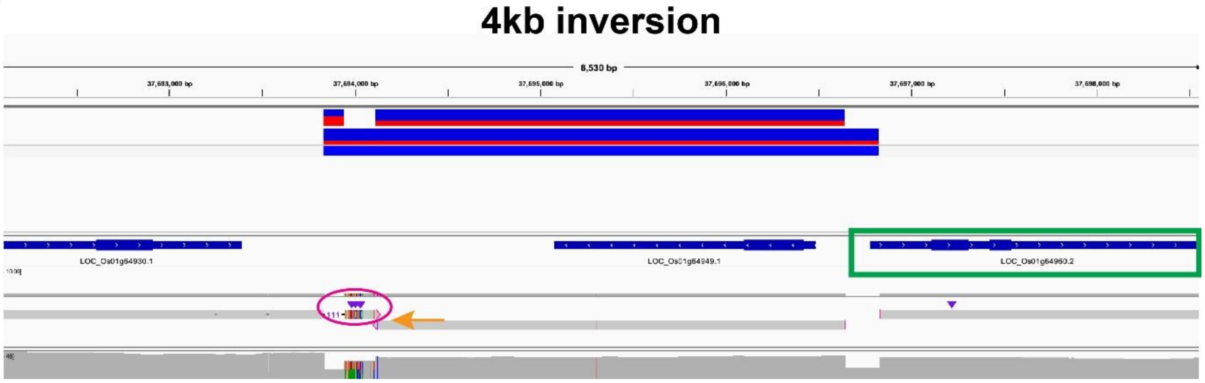


***Supplementary Figure 6.*** *High resolution view of the 19-1_OX inversion.*

**(a)** Increased resolution of the Chr 1 locus (~37.85 Mbp). **(b)** Integrative Genomics Viewer (IGV) display of raw mapped reads. The green box denotes the *OsPSBS1* gene (LOC_Os01g64960). Structural variant called by Sniffles is shown by the red and blue bar (Strength: 0.361). Evidence of the inversion is substantiated by head-to-head sequenced fragments (bottommost orange arrow). A non-reference, repetitive element is circled in magenta.

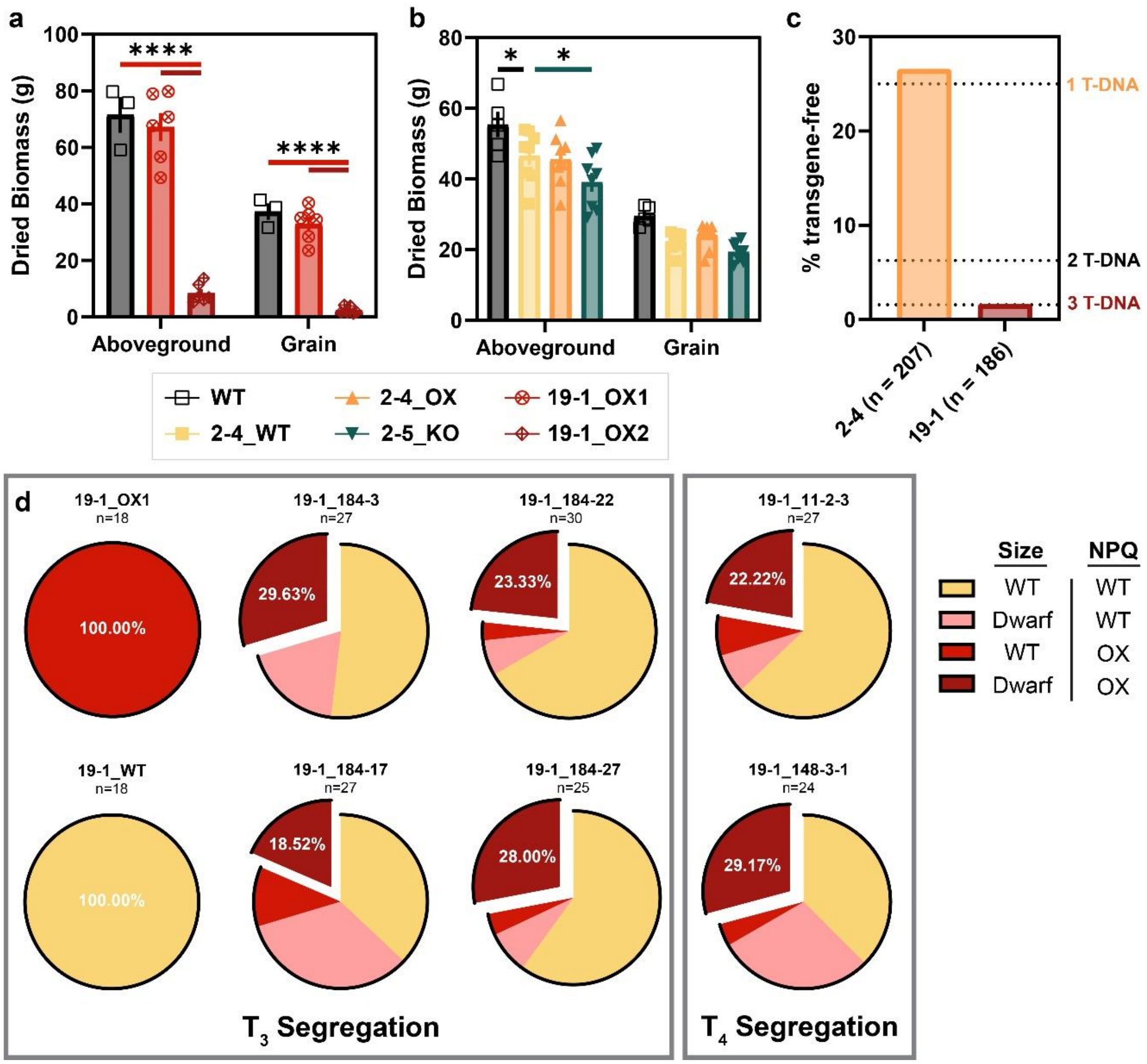


***Supplementary Figure 7.*** *Compromised biomass and non-mendelian inheritance of 19-1_OX2*

Differences in dried aboveground and grain biomass for selected **(a)** Event 19-1 and **(b)** Event 2 alleles. Genotypes are shown: WT (black, open square), 19-1_OX1 (red, crossed circle), 19-1_OX2 (maroon, crossed diamond), 2-5_KO (dark teal, inverse triangle), 2-4_WT (yellow, square), and 2-4_OX (orange, triangle). Pairwise significance in (a,b) was determined by ordinary one-way ANOVA (α=0.05) using Dunnett's test for multiple comparisons against Nipponbare WT or 2-4_WT, denoted by asterisks (*p≤0.05, ****p<0.0001). **(c)** Proportion of $T_1$ progeny sensitive to hygromycin and lacking the Cas9 transgene. **(d)** Percent inheritance of the parent allele as determined by biomass and NPQ phenotype across several generations. Alleles corresponding to 19-1_WT, 19-1_OX1, and 19-1_OX2 are shown in yellow, red, and maroon, respectively. Progeny dwarfed in size with WT NPQ are shown in pink.

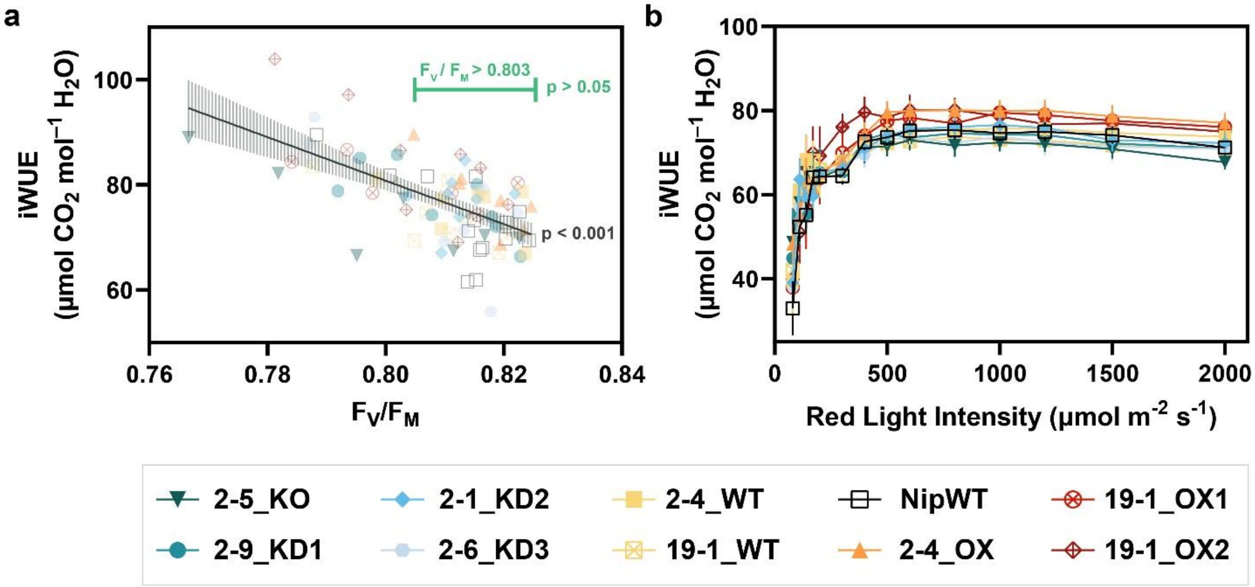


***Supplementary Figure 8.*** *Correlation of $F_v/F_m$ and iWUE across all genotypes.*

**(a)** Linear regression and 95% confidence interval of the average iWUE (µmol $CO_2$ $mol^{-1}$ $H_2O$) across all genotypes at light intensities ≥ 500 µmol $m^{-2}$ $s^{-1}$. Green bars mark the phenotypic range where the genotype-independent correlation between $F_v/F_m$ and iWUE is no longer significantly non-linear. **(b)** iWUE as a function of incident red light on mature flag leaves after constraining analysis to replicates $F_v/F_m > 0.803$ (n=4-7 biological replicates, data shown ± SEM). Genotypes are shown: WT (black, open square), 2-5_KO (dark teal, inverse triangle), 2-9_KD1 (teal, circle), 2-1_KD2 (blue, diamond), 2-6_KD3 (light blue, hexagon), 2-4_WT (yellow, square), 19-1_WT (yellow, crossed square), 2-4_OX (orange, triangle), 19-1_OX1 (red, crossed circle), 19-1_OX2 (maroon, crossed diamond).

***Supplementary Table 1.*** *OsPsbS1 non-coding sequence gRNA spacer sequences and positions relative to the start codon.*

| Insert | Orientation (rel. to ORF) | Position from ATG in *O. sativa sp. indica* cultivar IR64 | Position from ATG in *O. sativa sp. japonica* cultivar Nipponbare | Spacer Sequence (5' -> 3') |
|---|---|---|---|---|
| gRNA1 | R | -1163 : -1183 | -3837 : -3857 | GCGAGACACTAAAATACATT |
| gRNA2 | F | -973 : -953 | -3647 : -3627 | TCTTGTTCCTGGATGTAATT |
| gRNA3 | R | -888 : -908 | -3562 : -3582 | AGATTCAGGAGTAACAAAAA |
| gRNA4 | F | -769 : -749 | -3443 : -3423 | TGTTGCATGTGGTCCGTCGA |
| gRNA5 | R | -585 : -605 | -3259 : -3279 | CACAAAAAAGTACGGGAATG |
| gRNA6 | F | -374 : -394 | -374 : -394 | TACCAACCACCTGCTCTTCT |
| gRNA7 | R | -263 : -283 | -263 : -283 | GTCTCCCCGAATCCCTTCTA |
| gRNA8 | F | -166 : -146 | -166 : -146 | CTACGCCTCCCACCCGCCAC |
| gRNA scaffold | GTTTTAGAGCTAGAAATAGCAAGTTAAAATAAGGCTAGTCCGTTATCAACTTGAAAAAGTGGCACCGAGTCGGTGC | | | |
| tRNA linker | AACAAAGCACCAGTGGTCTAGTGGTAGAATAGTACCCTGCCACGGTACAGACCCGGGTTCGATTCCCGGCTGGTGCA | | | |

***Supplementary Table 2.*** *qRT-PCR primer pairs and observed primer efficiency in WT across a 1:81 dilution series.*

| Gene ID | Primer Name | Sequence | Primer Efficiency |
|---|---|---|---|
| UBQ[25] LOC_Os03g13170 | oDP578q UBQ-F_Fu2020 | TGCACCCTAGGGCTGTCAAC | 95.3% |
| | oDP579q UBQ-R-Fu2020 | GGCGAGTGACGCTCTAGTTCTT | |
| UBQ5[26] LOC_Os01g22490 | oDP580q UBQ5-F_Jain2006 | ACCACTTCGACCGCCACTACT | 105.1% |
| | oDP581q UBQ5-R_Jain2006 | ACGCCTAAGCCTGCTGGTT | |
| PsbS1[25] LOC_Os01g64960 | oDP584q PsbS1F-Fu2020 | CTGTTCGGCAGGTCCAAG | 95.9% |
| | oDP586q PsbS1R2-DP | CAAACCCGAGCATGGCGA | |

***Supplementary Table 3.*** *Primers used for PCR amplification and genotyping of OsPsbS1 NCS.*

| Amplicon | Forward Primer (5' -> 3') | Reverse Primer (5' -> 3') | Amplicon Size | Additional Sequencing Primer |
|---|---|---|---|---|
| Distal end (gRNA 1-5) | oDP214: AGACAGAGGTATGTCAATGTGTTATTG | oDP498: AAAGAGCAAATGGCCTACCA | 2384bp | oDP500: CGTGTCTCCCACGTCTTCTT |
| Proximal end (gRNA 6-8) | oDP497: TGGTAGGCCATTTGCTCTTT | oDP496: CATCCATCCAAATTCCAACCT | 2167bp | oDP499: GAGCAAACACTCAGGCACAA |
